# Comparing the stability and reproducibility of brain-behaviour relationships found using Canonical Correlation Analysis and Partial Least Squares within the ABCD Sample

**DOI:** 10.1101/2023.03.08.531763

**Authors:** Hajer Nakua, Ju-Chi Yu, Hervé Abdi, Colin Hawco, Aristotle Voineskos, Sean Hill, Meng-Chuan Lai, Anne L. Wheeler, Anthony Randal McIntosh, Stephanie H. Ameis

## Abstract

**Introduction:** Canonical Correlation Analysis (CCA) and Partial Least Squares Correlation (PLS) detect associations between two data matrices based on computing a linear combination between the two matrices (called latent variables; LVs). These LVs maximize correlation (CCA) and covariance (PLS). These different maximization criteria may render one approach more stable and reproducible than the other when working with brain and behavioural data at the population-level. This study compared the LVs which emerged from CCA and PLS analyses of brain-behaviour relationships from the Adolescent Brain Cognitive Development (ABCD) dataset and examined their stability and reproducibility.

**Methods:** Structural T1-weighted imaging and behavioural data were accessed from the baseline Adolescent Brain Cognitive Development dataset (*N* > 9000, ages = 9-11 years). The brain matrix consisted of cortical thickness estimates in different cortical regions. The behavioural matrix consisted of 11 subscale scores from the parent-reported Child Behavioral Checklist (CBCL) or 7 cognitive performance measures from the NIH Toolbox. CCA and PLS models were separately applied to the brain-CBCL analysis and brain-cognition analysis. A permutation test was used to assess whether identified LVs were statistically significant. A series of resampling statistical methods were used to assess stability and reproducibility of the LVs.

**Results:** When examining the relationship between cortical thickness and CBCL scores, the first LV was found to be significant across both CCA and PLS models (singular value: CCA = .13, PLS = .39, *p* < .001). LV_1_ from the CCA model found that covariation of CBCL scores was linked to covariation of cortical thickness. LV_1_ from the PLS model identified decreased cortical thickness linked to lower CBCL scores. There was limited evidence of stability or reproducibility of LV_1_ for both CCA and PLS. When examining the relationship between cortical thickness and cognitive performance, there were 6 significant LVs for both CCA and PLS (*p* < .01). The first LV showed similar relationships between CCA and PLS and was found to be stable and reproducible (singular value: CCA = .21, PLS = .43, *p* < .001).

**Conclusion:** CCA and PLS identify different brain-behaviour relationships with limited stability and reproducibility when examining the relationship between cortical thickness and parent-reported behavioural measures. However, both methods identified relatively similar brain-behaviour relationships that were stable and reproducible when examining the relationship between cortical thickness and cognitive performance. The results of the current study suggest that stability and reproducibility of brain-behaviour relationships identified by CCA and PLS are influenced by characteristics of the analyzed sample and the included behavioural measurements when applied to a large pediatric dataset.

## Introduction

Cortical structure has been linked to mental health symptoms across clinical and non-clinical samples (Romer et al., 2021; Ameis et al., 2014; Albaugh et al., 2016; Ducharme et al., 2016; Qin et al., 2014; Jacobs et al., 2020; Mihalik et al., 2019). Improving brain-behaviour research methods can lead to better characterization of risk or resilience to mental health symptoms (Nour, 2022). Yet, brain-behaviour associations have shown poor replicability across studies (Lombardo et al., 2019; Masouleh et al., 2019; Boekel et al., 2015), with recent work suggesting thousands of participants are required for replicable findings in cross-sectional studies (Marek et al., 2020). Small sample sizes, differences in between-study methodology, sampling heterogeneity, and underpowered statistical approaches have contributed to the lack of consistency between studies (Lombardo et al., 2019; Grady et al., 2021; Poldrack et al., 2017; Button et al., 2013). Recent multi-site research initiatives such as the Adolescent Brain Cognitive Development (ABCD) cohort study have largely addressed prior sample size limitations.

However, substantial methodological challenges remain, necessitating studies that can guide decisions around which statistical approaches are most suitable for identifying stable and reproducible brain-behaviour associations across studies. Brain-behaviour relationships have been often explored using univariate statistical approaches that assess one dependent variable per model, despite the limited power and sensitivity of this method when applied to complex brain-behaviour data (McIntosh, 2021; Mardia et al., 1979; Nakua et al., 2022). In contrast, multivariate approaches can examine relationships between several independent and dependent variables in a single analysis without requiring multiple comparison corrections, providing greater power to detect relationships between two variable sets (McIntosh, 2021; Marek et al., 2020).

Two widely used multivariate approaches to examine the association between two sets of variables are Canonical Correlations Analysis (CCA) and Partial Least Squares correlation (PLS). These approaches derive a set of latent variables (LV) pairs which each obtained as a linear combination of the variables of one data table. The latent variables in a pair have maximal correlation (using CCA) or covariance (using PLS) (Hotelling et al., 1936; Kotz & Johnson, 1985). This difference in the maximization criteria may result in the identification of different brain-behaviour relationships. Both CCA and PLS have been widely applied to clinical and population-based samples over the past decade to analyze brain metrics and phenotypic measures (Seok et al., 2021; Xia et al., 2018; Itahashi et al., 2020; Modabbernia et al., 2021; Ziegler et al., 2013; Kebets et al., 2019; Drysdale et al. 2017; Moser et al. 2018; Avants et al. 2014; Ing et al. 2019; Wang et al. 2018; Smith et al. 2015; Mihalik et al. 2019). As the accessibility of large-sample multidimensional datasets increases, CCA and PLS will likely be used more frequently given the potential to identify complex associations between multiple modalities.

Two prior studies have systematically compared the outputs from CCA and PLS models to determine the strengths, weaknesses, and similarities of both approaches (McIntosh, 2021; Helmer et al. 2020). Helmer et al. (2020) found that an adequate sample size (*N* > 1000) is necessary for stable and reliable CCA and PLS findings, regardless of the datatypes being examined. Although CCA and PLS both maximize linear relationships, McIntosh (2021) found that the identified relationships are most similar between CCA and PLS when the correlations within each data matrix are low. Beyond these findings, knowledge is limited regarding whether other characteristics of a dataset (e.g., measurements used, clinical versus non-clinical samples, etc.) influences the reliability of the models derived from CCA or from PLS. To address this gap, we applied CCA and PLS to brain and phenotypic behaviour data available through the ABCD dataset to: 1) examine the similarity between the identified LVs of CCA and PLS when delineating relationships between cortical thickness and phenotypic measures (which we refer to here as between-method generalizability), and 2) compare the stability, reproducibility, and reliability of the LVs within CCA and PLS (which we refer to here as within-method generalizability). We examined two different phenotypic measures to assess whether phenotypic characteristics influence brain-behaviour relationship identification when implementing CCA and PLS. The first analysis included the Child Behavior Checklist (CBCL)—a phenotypic measure of parent-reported childhood behaviours that are relevant to various mental health diagnoses. The second analysis examined relationships between cortical thickness and cognitive performance measured using the NIH Cognitive Toolbox. Using these two sets of measures provide an opportunity to compare whether the stability and reliability of CCA and PLS is influenced by the choice of measurement (i.e., parent-reported or performance-based) and the construct of interest (i.e., psychopathology or cognitive performance). The ultimate goal of our analyses is to determine whether CCA or PLS models are impacted differently by the various measures, and which approach may be better suited to identifying stable and reliable brain-behaviour associations in large population-based samples.

## Methods

### Sample

The ABCD dataset is a longitudinal multi-site population-based sample collecting a comprehensive measurement battery (including genetic, blood, environmental, cognitive, brain, and behavioural measures) in > 11,000 beginning in children who are 9-11 years old. Data collection time points occurring annually or biannually for 10 years. Participants were recruited from 21 academic sites across the U.S. using probability sampling to ensure that demographic trends across the U.S. are well represented in the sample (see Casey et al., 2018, for more details). Recruitment occurred through presentations and emails delivered to parents of children in local schools around each site. Interested parents underwent a telephone screening to determine whether their children were eligible to participate in the study. Participants were excluded from inclusion to the ABCD study if they had MRI contraindications, no English fluency, uncorrected vision or hearing impairments, major neurological disorders, were born extremely preterm (less than 28 weeks gestation), low birth weight (< 1200 grams), birth complications, or unwillingness to complete assessments. The current study used tabulated data from the baseline sample provided by the ABCD consortium from the fourth annual release (DOI:10.15154/1523041). Following removal of participants with poor brain imaging quality, missing T1-weighted scans, missing behavioural data, and removal of one sibling per family if the family enrolled multiple siblings, twins, or triplets (see *Figure S1* for consort diagram and details of exclusion), the current study used data from 9191 participants for the first analysis exploring associations between cortical thickness and CBCL scores (ages 9-11 years, 4369 assigned-female-at-birth and 4822 assigned-male-at-birth) and 9034 participants for the second analysis exploring associations between cortical thickness and NIH Toolbox scores (4292 assigned-female-at-birth and 4742 assigned-male-at-birth) .

### Scanning Acquisition and Processing

The neuroimaging protocol and specific T1-weighted parameters are detailed in previous publications (Casey et al. 2018; Hagler et al. 2018). Briefly, the ABCD protocol is harmonized for Siemens, General Electric, and Philips 3T scanners. All scanners used multi-channel coils capable of multiband echo planar imaging (EPI) acquisitions. The scanning occurred in either 1 or 2 sessions depending on the scanning site. Participants underwent a mock scanning session before the actual scan to help them get accustomed to the scanning environment. T1-weighted scans were collected, processed, and analyzed by the Data Analysis, Informatics Resource Center (DAIRC) based on standardized ABCD protocols (see details: Hagler et al. 2018).

Cortical and subcortical segmentation was performed using FreeSurfer v5.3.0 (Dale et al., 1999; Fischl and Dale, 2000). All T1-weighted scans were examined by trained visual raters who rated each scan from 0-3 based on motion, intensity homogeneity, white matter underestimation, pial overestimation, and visual artifacts. From these ratings, participants with poor quality scans were recommended for exclusion by the DAIRC and were excluded from this study (for more details, Hagler et al. 2018).

### Brain Measures

Cortical thickness of the 68 cortical regions from the Desikan-Killiany Atlas parcellations (Desikan et al., 2006) were used as the structural morphology measures in the current study.

### Behavioural Measures

Two different behavioural measures were selected for this analysis: CBCL subscale scores and the NIH Cognitive Toolbox. Both measures have been linked to structural brain morphology (Burgaleta et al., 2014; Ehrlich et al., 2012; Ronan et al., 2020; Ameis et al. 2014; Ducharme et al. 2014; Albaugh et al. 2016). The CBCL provides standardized parent-reported measures of behavioural symptoms relevant to mental health diagnoses (e.g., internalizing symptoms relevant to anxiety and depressive disorders), while the NIH Cognitive Toolbox offers a set of standardized performance-based measures of cognitive phenotypes (e.g., a working memory task).

In our first analysis, we implemented CBCL as the behavioural variable set. The CBCL is a well-validated tool to assess mental health symptoms in children from a parent/caregiver (Achenbach & Ruffle, 2000). The CBCL consists of 113 questions about a child’s behaviour on an ordinal scale (0 = never, 1 = sometimes, 2 = often). The ordinal ratings are summed to provide subscale scores for a variety of behaviours (e.g., aggression). We used the 8 conventional subscale scores from the parent-report CBCL for 6–18 year-olds: anxious/depressed, withdrawn/depressed, somatic complaints, thought problems, social problems, rule-breaking behaviour, aggressive behaviour, attention problems, in addition to 3 subscales included in ABCD that address symptoms closely related to neurodevelopmental disorders: stress symptoms, obsessive compulsive problems, and sluggish-cognitive-tempo (Jacobson et al. 2012; Storch et al. 2006).

In our second analysis, we used cognitive performance measures from the NIH Cognitive Toolbox as the behavioural variable set. The NIH Toolbox consists of 7 tasks presented on an iPad and measures 5 broad cognitive domains: picture vocabulary task (language skills), oral reading recognition task (language skills), list sorting working memory task (working memory), picture sequence memory task (episodic memory), flanker task (attention/inhibition), dimensional card change sort task (cognitive flexibility), and pattern comparison processing speed task (visual processing) (Thompson et al. 2019; Weintraub et al. 2013).

### Statistical Analysis

In order to remove the effects of age, sex-assigned-at-birth, total head size, site (*N* = 21), and MRI scanner model (*N* = 5) from the results, we regressed out (i.e., partialled out) these variables from both the brain and behavioural matrix using linear regression, a procedure ensuring that their effects would not drive the results. The resulting residuals were *Z*-transformed (mean-centered and standard deviation of 1) and were used in the CCA and PLS analyses.

#### Performing the CCA and PLS Analyses

CCA and PLS are unsupervised learning algorithms that identify linear relationships between two sets of variables, which describe the maximum relationship between 2 matrices (here a brain matrix, **X,** and a behaviour matrix, **Y**). These methods maximize different information between **X** and **Y** by decomposing a cross-product matrix. Usually, in CCA and PLS, the variables (i.e., columns) in **X** and **Y** are mean centered with unit variance (standard deviation of 1) or have a Euclidean norm of 1 (i.e., square root of the sum of squares equal to one). This scaling indicates that the cross-product matrix decomposed by PLS is either proportional (*Z-*scores are equal (norm 1)) to the Pearson correlation matrix between **X** and **Y** (denoted by **R_XY_**) and, the cross-product matrix decomposed by CCA is the *adjusted* Pearson correlation matrix between **X** and **Y** (denoted by **Ω**) as it is normalized by the within-block correlations of **X** and of **Y**. This difference in adjustment of within-block correlation makes the relationships derived from CCA optimized for correlation and those derived from PLS optimized for covariance (see details in supplementary Section 1, and in prior work: McIntosh et al. 2021; Krishnan et al. 2011; Abdi et al. 2018).

CCA and PLS maximize the associations between two data matrices by decomposing the **R_XY_** or **Ω** matrix with the singular value decomposition (SVD). The SVD decomposes **R_XY_** or **Ω** (for PLS and CCA, respectively) into 2 matrices of orthonormal singular vectors (**U** and **V**) and a diagonal singular value matrix (**S**). The singular vectors **U** and **V** store the **X** and the **Y** *loadings* that characterize the association between the two data matrices, respectively. The relationship between respective **U** and **V** singular vectors comprises *latent variables* (LVs; analogous to components, factors, or dimensions in the multivariate literature; Krishnan et al. 2011). The loadings give the weight of each variable in the corresponding LV (i.e., the degree to which that variable contributes to the latent relationship). For PLS, the loadings are directly derived from the **U** and **V** matrices, whereas for CCA, they are reweighted (see supplementary section 1 for details). To better understand the latent relationships between **X** and **Y**, each matrix is projected onto the respective singular vectors to create *latent scores* (i.e., **XU** and **YV**; analogous to factor scores). For each LV, the **XU-YV** pair of scores are optimized for correlation (in CCA) and for covariance (in PLS) to describe the multivariate co-variation between the **X** and the **Y** matrices. The singular values (from the **S** matrix) give the strength of the relationships between the corresponding pairs of latent scores from **XU** and **YV**. The largest possible number of LVs generated is equivalent to the minimum number of variables from either of the two matrices (e.g., when using CBCL as the behavioural measures, there were 11 LVs identified given that the behaviour matrix has 11 variables and the brain matrix has 68 variables).

The structure and distribution of **X** or **Y** may require some pre-processing prior to decomposing the cross-product matrix. In the current sample, the CBCL data were skewed with a high proportion of the sample having zero or low scores across subscales. As a result, cross-product matrices (**R_XY_** and **Ω**) were implemented using Spearman’s correlation when performing the CCA/PLS analysis using CBCL and NIH Toolbox scores to ensure methods remain consistent. Non-parametric Spearman’s correlation is more robust to skewed data (de Winter et al. 2016; Myers & Sirios, 2006).

To assess the significance of the LVs identified from CCA and PLS, we performed a permutation test on the singular values (McIntosh and Lobaugh, 2004; McIntosh, 2021). Briefly, the brain matrix underwent 10,000 iterations of resampling without replacement. Then, each resampled brain matrix and original behavioural matrix underwent the CCA or PLS analytical pipeline. Because the resampling procedure breaks the associations between the brain and behaviour matrices, this permutation creates the cross-product matrix under the null. Given the large sample size, a standard permutation test may not be stringent enough to detect meaningful LVs. To impose stringency on the detection of significant LVs, we performed a “sum of squares” permutation test (analogous to Wilk’s Lambda). This test assesses the *eigenspectrum* (i.e., a set of eigenvalues which are the square of the singular values) from CCA or PLS to determine if the total variance explained by a given set of LVs is greater than chance. It works by generating the sum of squares for *K* – 1 singular values (with 1 ≤ *K* < 11 (i.e., LV_1_-LV_1_) then LV_2_-LV_11_, etc.).

This permutation test generates null distributions of a given eigenspectrum depicting the likelihood of brain-behaviour relationships captured by a given LV occurring at random. Each LV was significant if less than 5% of the permuted sum of square singular values (e.g., LV_1_-LV_11_ for LV_1_, LV_2_-LV_11_ for LV_2_, etc.) were equal or greater than the empirical sum of square singular values.

#### Assessing Reproducibility, Reliability, and Stability

Resampling statistics were used to assess reproducibility, reliability, and stability of the LVs identified. Reproducibility of singular vectors were assessed using split-half resampling. Reliability of singular values was assessed using train-test resampling (*Figure 1*). Stability of the elements (i.e., individual variables, e.g., anxiety/depression subscale score from the CBCL) within a single LV was assessed using bootstrap resampling of the singular vectors. The split-half and train-test resampling analyses were conducted using 10,000 iterations. The bootstrap resampling of singular vectors analysis was conducted using 1000 iterations due to computational constraints. Reproducible and reliable distributions were identified using a *Z*-test (i.e., mean divided by standard deviation of distribution). A *Z*-test magnitude >1.96 is associated with a *p-*value <.05, a pattern indicating that the distribution significantly differed from zero. Prior work using simulated data confirms that randomly permuted distributions rarely exceed a *Z*-score of 1.65 (which would match a *p*-value of .10), a configuration suggesting that this cut-off is appropriate to use to determine significantly reproducible distributions (McIntosh, 2021).

**Figure 1.**
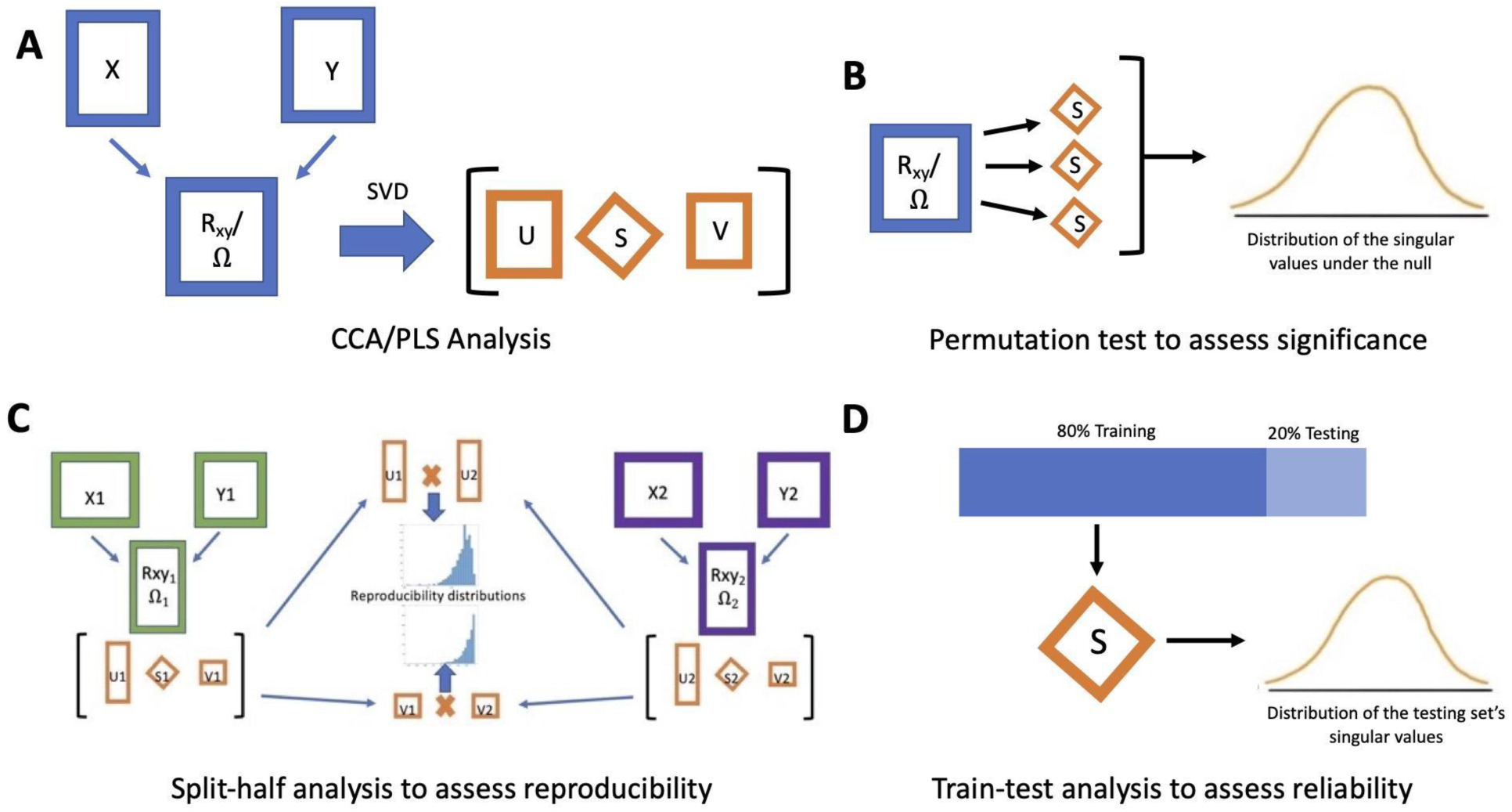
Overview of the analytical pipeline used in the current study. Panel A shows the main CCA and PLS analysis: the decomposition of the cross-product matrices to produce singular values and singular vectors (re-weighted in the case of CCA). Panel B shows the sum of squares permutation analysis to assess the statistical significance of each LV. Panel C illustrates the split-half analysis used to assess the similarity between respective singular vectors from each split-half. Panel D shows the train-test resampling analysis which assesses how well the singular values from the training sample can predict the singular values of the test sample. Note: interpretation of the different resampling statistical analyses is independent from one another and are not sequential. The bootstrap confidence interval estimation, not shown here, was used to assess the stability of the parameter estimates of variable weights in the singular vectors.

<u>Assessing reproducibility with split-half resampling:</u> For each iteration, the original **X** and **Y** matrices were randomly split into two halves to create 4 matrices (i.e., **X**_1_ and **Y**_1_ for the first half, and **X**_2_ and **Y**_2_ for the second half) conserving the relationship between **X** and **Y** for each resample (Churchill et al., 2013). CCA and PLS analysis were separately performed in each half (e.g., within a single iteration, CCA was conducted between **X**_1_ and **Y**_1_, and then separately between **X**_2_ and **Y**_2_). A Pearson correlation was used to assess the relationship between **U** and **V** singular vectors from the halves of each iteration (e.g., **U**_1_ and **U**_2_ from the analysis of the first and the second half). The *Z*-test was performed on the distribution of Pearson correlation values between each of the 11 singular vectors of matrices **U** and **V**.

<u>Assessing reliability with train-test resampling:</u> For each iteration, the data is split at random into an 80/20 train/test split. The CCA and PLS analyses were initially performed on the train set, and the singular vectors were projected onto the cross-product matrix of the test set to solve for the singular values (details in McIntosh, 2021). The *Z*-test was performed on the distribution of predicted singular values. This test assesses the magnitude of the association between **X** and **Y**.

<u>Assessing stability with bootstrap resampling:</u> Using a Monte-Carlo bootstrap approach, the data matrices **X** and **Y** were generated 1000 times by randomly selecting the participants with replacement until the total sample size was reached (*N* = 9191) while conserving the relationship between **X** and **Y**. The CCA and PLS were separately conducted for each regenerated **X** and **Y** matrix, a procedure resulting in 1000 regenerated matrices of singular vectors **U** and **V**. We corrected for arbitrary sign flips that often occur in iterations of the SVD (see supplementary section 1.2 for details). The 1000 regenerated **U** and **V** singular vectors provide *bootstrap samples* that were used to create the *bootstrapped distribution* of each element in **U** and **V**. From these bootstrapped distributions, we computed the 95% *bootstrapped* confidence intervals of each element in **U** and **V** to quantify their stability. A stable variable in a given LV was determined by a corresponding 95% bootstrapped confidence interval that does not include zero.

### Sensitivity analyses

We ran two sensitivity analyses to ensure that the differences in the CCA and PLS results were not driven by socioeconomic status (SES) or history of head injury, both of which have been found to impact structural morphology metrics and/or behaviour (Lawson et al. 2013; Piccolo et al. 2016; Wilde et al. 2012). For both these samples, we performed the CCA and PLS analysis, permutation test, and split-half resampling analysis. We analyzed a subset of the sample with available household income data (*N* = 8399) which we used as a proxy for SES (Hill et al., 2016; Rakesh et al., 2021). In this analysis, household income data were included as a covariate in the linear regression model to extract residuals from the brain and behaviour matrices. In a separate analysis, we performed CCA and PLS in a subset of the sample including participants with no history of head injuries (*N* = 8139; see details in supplementary section 3).

### Post-hoc analyses

Between- and within-method generalizability may be improved by stratifying the sample based on clinical severity when examining the relationship between cortical thickness and CBCL scores. We therefore implemented two *post-hoc* analyses to explore whether variations of the sample or measures improves the reproducibility of brain-behaviour relationships derived from CCA or PLS. Given the skewed nature of CBCL scores, we sought to determine whether participants with higher scores (i.e., greater behavioural problems) would have specific brain-behaviour relationships that may be washed out when conducting the analysis in the full sample. We stratified the sample by participants who had a total CBCL T-score (normalized for sex-assigned-at-birth and age) >60 which has been proposed as a subclinical cut-off (*N* = 1016; (Biederman et al., 2020)). In a separate post-hoc analysis, in order to remove the zero inflation (Byrd et al. 2021) we removed participants with no endorsement (i.e., a score of 0) on any CBCL subscale to attain a sample with full subscale symptom endorsement (*N* = 5196; see supplementary section 4). We performed the CCA and PLS analysis, permutation test, and split-half resampling analysis in both these subsets.

**Data and Code Availability:** Data for the ABCD Study are available through the National Institutes of Health Data Archive (NDA; nih.nda.gov). The participant IDs included in these analyses and details on the measures used, can be found in this project’s NDA study (DOI: 10.15154/1528644). The code for the analysis can be found on GitHub (https://github.com/hajernakua/cca_pls_comparison_ABCD).

## Results

### Participant Characteristics

*Table 1* provides the demographic details of the ABCD sample used in the current study. As shown, there were no substantial differences between the sex, household income, race/ethnicity, parental education, and behaviour measures between the analyzed sample (*N* = 9191) and the total sample acquired from ABCD with complete data (*N* = 11804). See *Table S1* comparing diagnostic characteristics between the sample with full CBCL data (*N* = 9191) and analyzed subsamples (*N* = 8399, 8139) included in sensitivity analyses (see supplementary section 3 for details on sensitivity analyses).

**Table 1.** Demographic Characteristics of the ABCD subsample included in the current study and the acquired ABCD sample

|  | Included ABCD<br>Sample (n=9191) |  | Acquired ABCD<br>Sample (n=11804) |  |
| --- | --- | --- | --- | --- |
|  | Mean [range] | SD | Mean [range] | SD |
| Age (in months) | 118.9 [107-133] | 7.4 | 118.9 [107-133] | 7.5 |
| CBCL Total T-Score | 46.12 [24-83] | 11.4 | 45.84 [24-83] | 11.3 |
| CBCL Internalizing T-Score | 48.73 [33-93] | 10.7 | 48.45 [33-93] | 10.64 |
| CBCL Externalizing T-Score | 45.87 [33-84] | 10.4 | 45.73 [33-84] | 10.32 |
|  | **Total** | **%** | **Total** | **%** |
| Sex (Female) | 4369 | 47.5 | 5637 | 47.7 |
| *Household Income* |  |  |  |  |
| <\$50K | 2531 | 27.5 | 3196 | 27.1 |
| \$50-\$100K | 2381 | 25.9 | 3057 | 25.9 |
| >\$100K | 3487 | 37.9 | 4544 | 38.5 |
| *Participant race/ethnicity* |  |  |  |  |
| White | 4704 | 51.2 | 6150 | 52.1 |
| Black | 1360 | 14.8 | 1755 | 14.9 |
| Asian | 205 | 2.2 | 251 | 2.12 |
| Hispanic | 1973 | 21.5 | 2401 | 20.3 |
| Other | 948 | 10.3 | 1244 | 10.5 |
| *Parent Education* |  |  |  |  |
| <HS Diploma | 468 | 5.1 | 583 | 4.9 |
| HS Diploma | 887 | 9.65 | 1120 | 9.5 |
| Some College | 2389 | 25.9 | 3056 | 25.9 |
| Bachelors | 2287 | 24.9 | 3003 | 25.4 |
| Post-Graduate | 3150 | 34.3 | 4027 | 34.1 |

Note: Some participants had missing household income, race/ethnicity or parent education information, thus, the total number of participants in each category reflect those with available data. SD = standard deviation; CBCL = Child Behavior Checklist, HS=high school. Internalizing behaviour is the summed broad-band measure which includes anxious/depressed, withdrawn/depressed, and somatic complaints scores. Externalizing behaviour is the summed broad-band measure which includes aggressive and rule-breaking behaviour. These broad-band measures are clinically meaningful, however, they do not allow the exploration of brain-behaviour relationships at the symptom level. The second analysis included all the participants in the first with available NIH Cognitive Toolbox data (n=9034). There were limited differences in demographic characteristics between the sample with complete CBCL data and the sample with complete NIH data.

### CCA and PLS Analyses

*First Analysis (CBCL scores as the behavioural variable set):* the sum of squares permutation test revealed 1 significant LV for both CCA and PLS (*p* < .001) when decomposing the Spearman’s cross-product correlation matrix. The relationship between the brain (**U**) and behaviour singular vectors (**V**) in LV_1_ was stronger in PLS compared to CCA (singular values: PLS = .39 [81.6% of covariance], CCA = .13 [19.3% of variance]). *Figure 2* depicts un-thresholded behaviour and brain loadings for PLS and CCA. LV_1_ from PLS identified an association between lower CBCL scores (i.e., less behavioural problems) and decreased cortical thickness; the strongest loading was between aggressive behaviours and the right pars triangularis. LV_1_ from CCA identified both positive and negative behaviour loadings linked to both increased and decreased cortical thickness; the strongest loading was between greater social problems and the greater thickness of the right superior temporal gyrus. See *Figure S2* for CCA and PLS results when decomposing the Pearson correlation matrix. See supplementary section 1.2 and *Figure S4* for CCA results when implementing the structure coefficients. See supplementary section 3 and *Figure S3* indicating similar results found for sensitivity analyses controlling for household income and head injuries in the sample.

**Figure 2.**
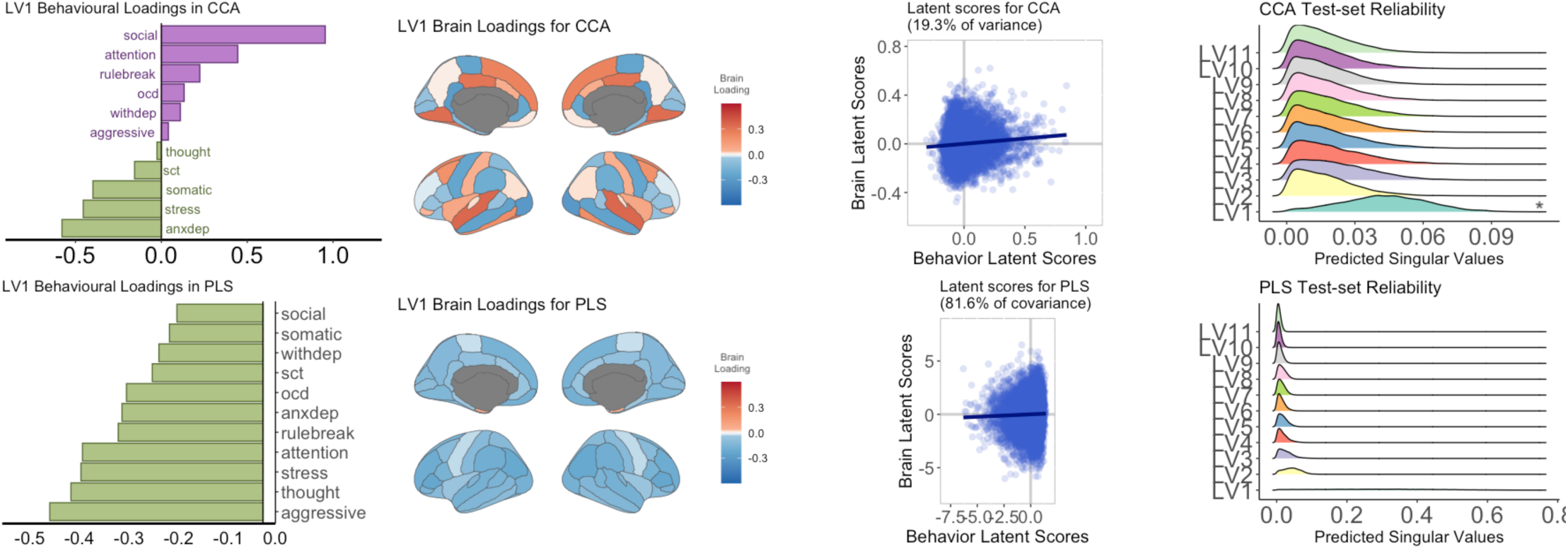
Unthresholded behaviour and brain loadings from the PLS and CCA analysis performed in the first analysis (n=9191). The highest behaviour loadings for PLS were aggressive behaviour, thought problems, and stress. The highest brain loadings for PLS were right pars triangularis, right inferior parietal cortex, and left posterior cingulate cortex. The highest behaviour loadings for CCA were social problems, withdrawn/depressive symptoms, and attention problems. The highest brain loadings for CCA were left and right posterior cingulate cortex and left fusiform gyrus. Importantly, interpretation of the loading direction is relative. As such, it is accurate to say that in PLS negative behavioural loadings are linked to negative brain loadings or that positive behavioural loadings are linked to positive brain loadings as long as the relationship of the sign remains the same. The second farthest right column shows the latent scores between *XU* and *YV* for LV1. Prior to calculating the latent scores, the brain and behavioural loadings have been standardized by the singular values. The *x*-axis from the train-test distributions is the predicted singular values of the test sample for each iteration. Astericks indicate the LVs which showed a distribution with a Z-score greater than 1.96 LV1 from CCA was found to be reliable (that is, LV1 of the training sample can reliably predict the singular values of LV1 from the test sample). OCD = obsessive compulsive symptoms, withdep = withdrawn/depression symptoms, sct = sluggish-cognitive-tempo, anxdep = anxiety/depression symptoms, rulebreak = rule breaking behaviour.

*Second Analysis (NIH Toolbox scores as the behavioural variable set):* the permutation test revealed 6 significant LVs (LV_1_: PLS = .43 [75.5% of variance]; CCA = .21 [41.6% of variance]; *Figure 3*). In contrast to the first analysis, LV_1_ between cortical thickness and performance on the NIH cognitive toolbox were similar in CCA and in PLS, indicating the presence of between-method generalizability. In both CCA and PLS, this LV identified an association between decreased cognitive performance (across all variables) and increased cortical thickness in several frontal and temporal regions and decreased cortical thickness in several occipital regions (*Figure 3*). The strongest associations in LV_1_ were between the list sorting working memory task (working memory) and the picture vocabulary task (language abilities) and the left pars opercularis and left parahippocampal gyrus. See supplementary section 3 and *Figure S7* indicating similar results found for sensitivity analyses controlling for household income and head injuries in the sample.

**Figure 3.**
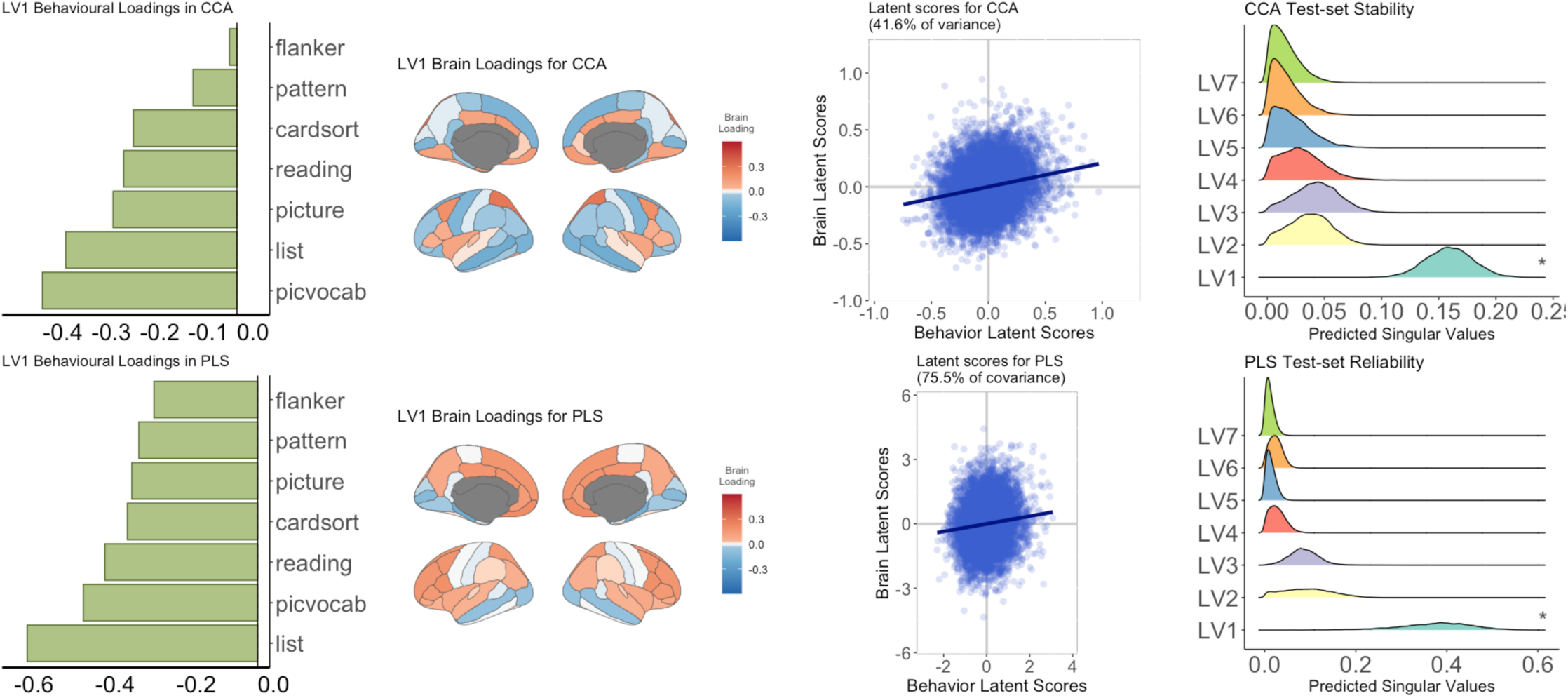
Unthresholded behaviour and brain weights from the PLS and CCA analysis performed between NIH Cognitive Toolbox scores and cortical thickness. The highest 2 behaviour loadings for PLS and CCA were performance on the list sorting task and the picture vocabulary task. The highest brain loadings for CCA were the left pars opercularis, superior frontal gyrus, and parahippocampal gyrus. The highest brain loadings for PLS were the left pars opercularis, parahippocampal gyrus, and medial orbitofrontal gyrus. The second farthest right column shows the latent scores between *XU* and *YV* for LV1. Prior to calculating the latent scores, the brain and behavioural loadings have been standardized by the singular values. Overall, there is a similar relationship between the brain and behavioural latent scores when comparing CCA and PLS. The fourth (farthest right) column shows the results of the train-test resampling analysis. Astericks indicate the LVs which showed a distribution with a Z-score greater than 1.96. LV1 for both PLS and CCA was found to be reliable (that is, LV1 of the training sample can reliably predict the singular values of LV1 from the test sample). Flanker = Flanker Task, pattern = pattern comparison processing speed task, cardsort = dimensional change card sort task, reading = oral reading recognition task, picture = picture vocabulary task, list = list sorting working memory task, picvocab = picture vocabulary task.

## Reproducibility, Reliability, and Stability

*Split-half resampling (reproducibility of loadings): Figure 4* shows the distributions of Pearson correlation coefficients between the **U** (or **V**) singular vectors from the two halves. In the first analysis (CBCL scores as behavioural matrix), the *Z*-test of the distribution shows that the correlation between the loadings from both halves is not significantly different from zero across all LVs, indicating no reproducible singular vectors for CCA. A similar pattern was found in PLS except for the behaviour loadings for LV_2_ of which the mean coefficient of correlation is significantly different from 0 (*Z*-score = 2.17), a value indicating some reproducibility. In the second analysis (NIH Toolbox scores as behavioural matrix), LV_1_ for the behavioural and brain loadings was found to be reproducible for both CCA and PLS (Z-score range = 2.9 – 25). LV_3_ behavioural and brain loadings were found to be reproducible for PLS (Z-score = 2.15, 2.57, respectively).

**Figure 4.**
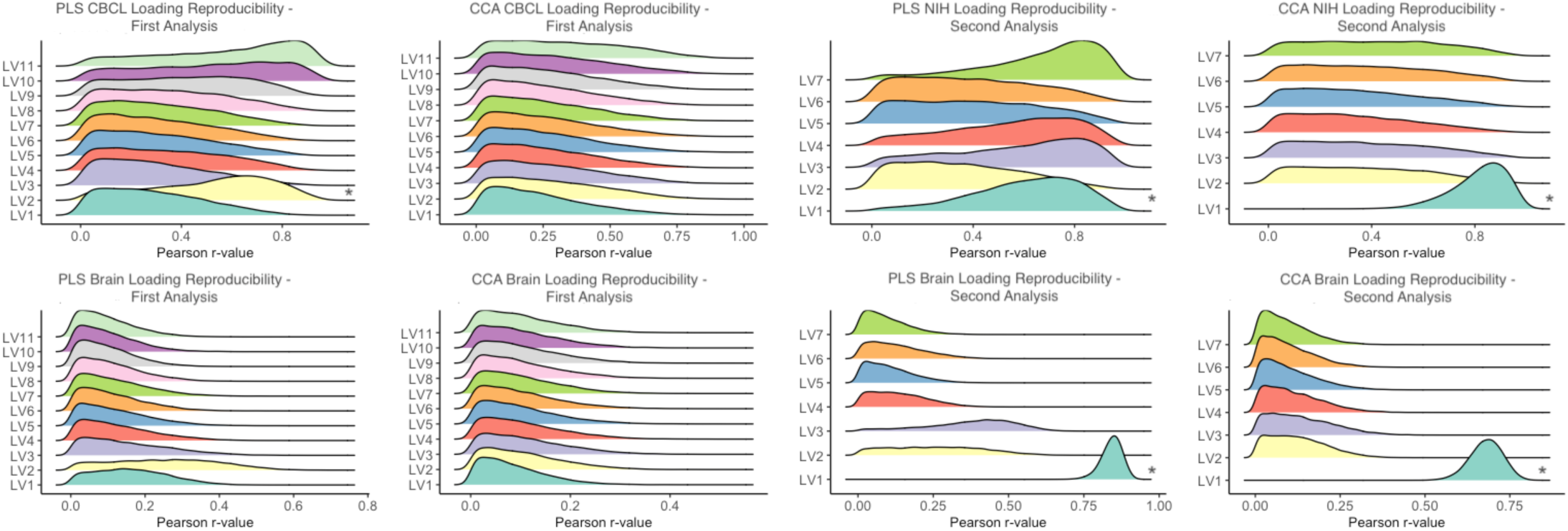
This figure depicts the distributions of the resampled loadings from the split-half analysis for the first and second analyses. The x-axis from the split-half distributions are the Pearson correlation coefficients between respective loadings from each split-half analysis (e.g., *U1* and *U2* from the analysis comparing X1 and Y1 and separately, X2 and Y2). Astericks indicate the LVs which showed a distribution with a Z-score greater than 1.96.

*Train-test resampling (reliability of singular values): Figure 2 and 3* (farthest right panel) illustrates the distributions of predicted singular values of each test set iteration. In the first analysis (CBCL scores as behavioural matrix), the results showed that LV_1_ from CCA features a distribution significantly different from zero (*Z*-score = 2.31), indicating a reliable LV (i.e., the SVD results from the training set can reliably predict the singular values of the test set). No other LV showed reliable singular values in CCA or PLS. For the second analysis (NIH Toolbox scores as behavioural matrix), LV_1_ and LV_3_ were found to be reliable for both CCA and PLS (Z-score range = 2.15 – 7.77), and LV_2_ was additionally found to be reliable for CCA (Z-score = 2.02).

*Bootstrap resampling (stability of elements in LV_1_):* Bootstrap analysis on the loadings from the first and second analyses revealed several stable brain and behavioural elements in LV_1_ (i.e., the 95% confidence intervals did not include zero; see *Figures S13-S14*).

## Post-hoc analyses

In the subset of the sample with elevated total CBCL scores (elevated-CBCL sample, T-score > 60; *N* = 1016), the correlations in the cross-product matrices **R_XY_** and **Ω** were overall stronger than those of the main sample (*Figure S11*). Accordingly, the first singular values were also larger than found in the main sample, a difference indicating a stronger relationship between the brain and behavioural variables (singular values of LV_1_: PLS = .67 [54% of covariance]; CCA = .33 [14.6% of variance]). The PLS analysis identified higher CBCL scores (i.e., greater problems) linked to lower cortical thickness across nearly all regions, the opposite finding to the main analysis using CBCL scores as the behavioural matrix (*Figure S5*). However, none of the LVs extracted from the elevated-CBCL sample from both PLS and CCA were statistically significant or reproducible. By contrast, from the subset excluding participants with full CBCL subscale endorsement (*N* = 5196), the relationships between brain and behaviour variables were similar to that of the main sample but were not statistically significant or reproducible (singular values of LV_1_: PLS = .38, CCA = .18; *Figure S6*).

## Discussion

A challenge in clinical neuroscience research is identifying brain-behaviour relationships that are consistent across different samples and studies. Here, we aimed to better understand factors that influence between-method (i.e., CCA vs. PLS) and within-method (i.e., reproducibility/reliability/stability) generalizability of brain-behaviour relationships identified using CCA and PLS applied to cortical thickness and behavioural data from the baseline ABCD sample. In our first analysis using CBCL scores, the results indicated that LV_1_ identified different multivariate patterns of brain-behaviour associations using CCA or PLS, a difference suggestive of limited between-method generalizability. Neither CCA nor PLS LVs were found to be consistently stable or reproducible when systematically examined through various resampling statistical approaches, suggestive of low within-method generalizability. In our second analysis using cognitive performance measures, LV_1_ identified similar brain-behaviour relationships using both CCA and PLS which were reproducible, reliable, and stable suggestive of both between- and within-method generalizability. Our findings suggest that using performance-based measures may be more optimal for delineating multivariate relationships that are generalizable (between- and within-method) between cortical thickness and phenotypic data in large community-derived samples, consistent with some prior work (Marek et al., 2020; Ooi et al., 2022).

## CCA and PLS identifies different multivariate relationships between cortical thickness and parent-reported behavioural measures (i.e., low between-method generalizability)

In our first analysis examining associations between cortical thickness and parent-reported behaviour using the CBCL, LV_1_ derived from CCA indicated that both higher and lower CBCL scores were linked to both higher and lower cortical thickness values. The pattern of covariation found in CCA is consistent with prior work examining brain-phenotype relationships in large community derived samples. One such study found covariation of brain and phenotypic measures such that positive and negative environmental factors loaded in opposite directions and were linked to covariation of functional connectivity (Smith et al. 2015). CCA may be more likely to identify prominent patterns of covariation (i.e., positive and negative loadings) between two data matrices due to the adjustment of the within-block correlation matrices in the cross-product matrix (**Ω**) submitted to the SVD. This step removes the redundant within-block variance optimizing for uniqueness.

LV_1_ derived from PLS, in contrast, identified lower CBCL scores (i.e., fewer problems) linked to mainly decreased cortical thickness. The limited covariation in LV_1_ (i.e., the various behavioural and brain variables load in the same direction) from PLS may be due to the optimization of redundant relationships given that the cross-product matrix does not adjust for within-block correlations. The limited covariation has been found in prior reports examining relationships between mental health symptoms and functional connectivity using PLS (Itahashi et al. 2020). Notably, the brain-behaviour relationship in LV_1_ for PLS in the main sample contrasts with prior univariate analyses reporting negative associations between CBCL measured behavioural problems and cortical thickness in clinically enriched or normative samples (Ameis et al., 2014; Ducharme et al., 2014, 2011; Jacobs et al., 2020; Vijayakumar et al., 2017).

However, in the elevated-CBCL sample (i.e., participants with CBCL total t-scores >60; *N* = 1016), we found that LV_1_ in PLS linked higher behavioural scores to lower cortical thickness. Although the results were not significant, the increased correlations in the cross-product matrix (*Figure S11*) suggest that a larger sample would be needed to find significant results. The contrast of these results suggest that brain-behaviour relationships identified at the population level may be washing out brain-behaviour patterns present in children with more pronounced behavioural problems. Overall, these findings indicate low between-method generalizability for multivariate relationships between parent-reported behavioural symptoms and cortical thickness.

## Lack of consistently stable and reproducible LVs linking cortical thickness and CBCL scores (i.e., low within-method generalizability)

In the first analysis, there was no consistent reproducible or reliable LVs linking cortical thickness and CBCL scores as assessed by split-half or train-test resampling, respectively. The LV_2_ behavioural loading was reproducible when implementing PLS. The singular value for LV_1_ in CCA was found to be reliable as assessed by train-test resampling, a result indicating a robust, albeit small, signal greater than random chance (McIntosh & Lobaugh, 2004). Further, there were stable CBCL and cortical thickness elements in LV_1_ as assessed by bootstrap resampling using PLS, but not with CCA. These findings suggest that tests of reproducibility, reliability, stability, and significance may not lead to consistent conclusions about the generalizability of brain-behaviour relationships as was found in prior reports (McIntosh, 2021). Notably, despite limitations in between- and within-generalizability, LV_1_ was robust (i.e., results remained similar despite changes to the sample, as noted by Huber, 2011) given that the sensitivity and post-hoc analysis in the subsample with full subscale endorsement did not substantially alter the relationship identified in LV_1_ for CCA or PLS. The statistical significance and robustness of LV_1_ indicates that the results are meaningful to the present sample; however, the lack of consistent reproducibility and reliability suggests that the results are not generalizable to other samples.

## High between- and within-method generalizability between cortical thickness and cognitive performance

In the second analysis examining associations between cortical thickness and cognitive performance from the NIH Cognitive Toolbox, LV_1_ derived from CCA and PLS linked cognitive performance scores to both higher and lower cortical thickness values. In contrast to the trends found when using parent-report CBCL as behavioural measures, CCA and PLS both found covariation in the brain loadings but not in the behavioural loadings. This difference may be due to the stronger between-block correlations in the cross-product matrix (**R_XY_** and **Ω**; *Figure S10*) compared to the first analysis leading the NIH scores to exhibit the same directional relationship to cortical thickness regardless of whether CCA or PLS was implemented.

Notably, the stability and reproducibility of LV_1_ across methods suggests that the results found are both meaningful to the current sample (given the statistical significance and robustness) and generalizable to other samples. Thus, between-method generalizability (i.e., similar LV_1_ between CCA and PLS) and within-method generalizability (i.e., consistent stability and reproducibility of LV_1_) suggest that performance-based measures may be better to use when exploring latent relationships between behavioural phenotypes and brain structural morphometry.

## Skewed data and low correlations may reduce between- and within-method generalizability

In the first analysis, the lack of between- and within-method generalizability found between the brain and parent-report behavioural variables may have been driven by the low correlations (i.e., minimal effect size) as well as the skew of the CBCL variables (*Figure S12*). The skewed CBCL scores indicate that the variance of interest is present within a small subset of the model with the most extreme behavioural problems. Importantly, low brain-behaviour correlations and the skewness of behaviour scores are likely a feature of population-based samples, not the result of sampling error (Owens et al. 2021). In contrast, the NIH Toolbox measures are more normally distributed (*Figure S12*) and feature slightly stronger correlations between the brain and behavioural variable sets (i.e., cross-correlation matrix, *Figure S10*). These configurations likely contributed to the increased between- and within-method generalizability when using cognitive performance measures in the current study. The differences in within- and between-method generalizability between the first and second analyses may also be due to the distribution of the variance in the NIH toolbox scores compared to CBCL scores. To better understand the variance structure of CBCL and NIH Toolbox scores, we submitted each of these two matrices to a separate principal components analysis. CBCL had a more compressed variance structure compared to NIH Toolbox scores (CBCL: 6/11 (54.5%) and NIH: 5/7 (71%) of the components were needed to explain 90% of the variance). This compressed variance structure may be another explanation contributing to the limited within-method generalizability of the PLS and CCA models.

Prior work has noted the limitations of clinical measures that are originally optimized to detect clinically significant symptoms which lead to skewed distributions when applied to population-based or community-derived samples (Alexander et al., 2020). More recent work has shed light on the importance of developing more rigorous phenotypic measures, particularly in population-based samples, as limited intraclass reliability has been shown to impact clinical neuroscience research (Nikolaidis et al. 2022). The results of the current study extend this prior work and provide evidence in support of either collecting larger samples of participants with greater clinical impairments and/or development of phenotypic measures that adequately capture the variation of behavioural presentations at the population level.

The results of the current study suggest that reproducibility, reliability, and stability of CCA and PLS are largely impacted by the variance structure of the phenotypic measure used when examining brain-behaviour relationships. Although we did not find major differences between CCA and PLS in the second analysis, these two approaches do have different underlying philosophies that may be important when considering which model to implement. The removal of the within-block correlations allows CCA to amplify the specific and unique variable(s) that most strongly categorizes the **Ω**. In contrast, PLS takes advantage of the redundancy of the within-block correlations and amplifies the collection of variables that most strongly characterizes **R_XY_**. Importantly, the one-to-one comparison between CCA and PLS was possible in this study due to the use of ROI data for both analyses. Given that CCA maximizes for correlations, very high multicollinearity within a variable set will result in unstable LVs with CCA (e.g., if using voxel-wise data). In such cases, PLS should be used.

## Limitations

There are some limitations to consider when interpreting the results of this study. First, we analyzed the baseline data from the ABCD Study which includes children between 9-11 years of age. This age range may not feature substantial variability of cortical thickness to facilitate strong relationships linked to parent-reported behavioural phenotypes. Second, we focused our analysis on classical CCA and PLS as opposed to more recent derivative approaches, such as kernel CCA or sparse PLS (Mihalik et al., 2019; Witten & Tihshirani, 2009). These derivative approaches add a penalty factor to the decomposition step, reducing the weight of variables that have weak contributions to the overall LV, thereby facilitating easier interpretability and increasing reproducibility of results. These approaches have typically been used in datasets which include a disproportionate ratio of participants to variables (i.e., low sample-to-feature ratio, Mihalik et al. 2019) which was not the case in the current sample. While comparing the derivative approaches is beyond the scope of the current paper, it is important to explore in future work.

## Conclusion

Clinical neuroscience research is going through a translational crisis largely due to the challenges of delineating brain-phenotype relationships that are replicable, meaningful and generalizable, particularly with respect to behaviours linked to mental health diagnoses (Nour et al. 2022). There have been methods that address this challenge, such as the use of an external validation sample to determine generalizability (Scheinost et al., 2019; Walter et al., 2019; Yip et al., 2020). Less emphasis has been placed on assessing generalizability of results when using different methods within the same sample. The results of the current study suggest that between- and within-method generalizability among CCA and PLS is influenced by sample/measurement characteristics. Low correlations between brain and behavioural measures coupled with skewed distribution reduces reproducibility, reliability, and stability of CCA and PLS models. The results of this study suggest that the measures inputted into CCA or PLS models play a more substantial role in the generalizability of the model’s results compared to the specific approach applied (i.e., CCA or PLS). One important implication of the current work is that performance-based cognitive measures are likely a more promising phenotypic measure compared to parent-reported behaviour in children when examining multivariate relationships with brain structure.

There are several avenues of future work that should be examined to better understand how to improve brain-behaviour relationship delineation at the population-level. First, it is possible that different sub-groups of participants may show differing relationships between brain structure and complex phenotypic outcomes such as parent-reported CBCL scores, which may reduce the stability of findings when considering a single linear dimension across the entire sample. As a result, biotyping/clustering methods may be better suited when delineating brain-behaviour relationships using clinical report measures. Second, the current findings are specific to cross-sectional investigations of cortical thickness and phenotypic measures. It is possible that using longitudinal phenotypic data, available per individual across different time-points, as part of the ABCD longitudinal data collection will reveal within-person stability and reproducibility of brain-behaviour relationships. Lastly, it is possible that using functional MRI acquisitions, which feature greater variability compared to structural MRI metrics, for the brain matrix, may result in greater between- and within-method generalizability when linked to parent-report behavioural measures.

**Funding, Disclosures, and conflict of interest:** HN has received funding from the CAMH Discovery Fund, Ontario Graduate Scholarship, Fulbright Canada, and currently receives funding from the Canadian Institutes of Health Research (CIHR) Doctoral Award. CH current receives funding from the National Institute of Mental Health (NIMH), The CAMH Foundation, Natural Sciences and Engineering Research Council of Canada (NSERC). ANV currently receives funding from the NIMH, CIHR, Canada Foundation for Innovation, CAMH Foundation, and University of Toronto. M-CL receives funding from CIHR, the Academic Scholars Award from the Department of Psychiatry, University of Toronto, and CAMH Foundation. ALW currently receives funding from CIHR, Brain Canada Foundation, and NSERC. ARM currently receives funding from NSERC and CIHR research grant. SHA currently receives funding from the NIMH, CIHR, the Academic Scholars Award from the Department of Psychiatry, University of Toronto, and the CAMH Foundation. Other authors report no related funding support, financial or potential conflicts of interest.

## Acknowledgements

Data used in the preparation of this article were obtained from the Adolescent Brain Cognitive Development® (ABCD) Study (https://abcdstudy.org), held in the NIMH Data Archive (NDA). This is a multisite, longitudinal study designed to recruit more than 10,000 children age 9–10 and follow them over 10 years into early adulthood. The ABCD Study® is supported by the National Institutes of Health and additional federal partners under award numbers U01DA041048, U01DA050989, U01DA051016, U01DA041022, U01DA051018, U01DA051037, U01DA050987, U01DA041174, U01DA041106, U01DA041117, U01DA041028, U01DA041134, U01DA050988, U01DA051039, U01DA041156, U01DA041025, U01DA041120, U01DA051038, U01DA041148, U01DA041093, U01DA041089, U24DA041123, U24DA041147. A full list of supporters is available at https://abcdstudy.org/federal-partners.html. A listing of participating sites and a complete listing of the study investigators can be found at https://abcdstudy.org/consortiummembers/. ABCD consortium investigators designed and implemented the study and/or provided data but did not necessarily participate in analysis or writing of this report. This manuscript reflects the views of the authors and may not reflect the opinions or views of the NIH or ABCD consortium investigators. The ABCD data repository grows and changes over time. The ABCD data used in this report came from NDA Release 4.0 (DOI:10.15154/1523041).

## References

Abdi, H., Guillemot, V., Eslami, A., & Beaton, D. 2018. Canonical Correlation Analysis. Encyclopedia of Social Network Analysis and Mining, doi 10.1007/978-1-4614-7163-9_110191-1

Achenbach, T.M., Ruffle, T.M., 2000. The child behavior checklist and related forms for assessing behavioral/emotional problems and competencies. Pediatr. Rev. 21, 265–271. https://doi.org/10.1542/pir.21-8-265

Albaugh, M.D., Ducharme, S., Watts, R., Lewis, J.D., 2016. Anxious / depressed symptoms are related to microstructural maturation of white matter in typically developing youths.

Alexander, L.M., Salum, G.A., Swanson, J.M., Milham, M.P., 2020. Measuring strengths and weaknesses in dimensional psychiatry. J Child Psychol Psychiatry 61, 40–50. https://doi.org/10.1111/jcpp.13104

Ameis, S.H., Ducharme, S., Albaugh, M.D., Hudziak, J.J., Botteron, K.N., Lepage, C., Zhao, L., Khundrakpam, B., Collins, D.L., Lerch, J.P., Wheeler, A., Schachar, R., Evans, A.C., Karama, S., 2014. Cortical thickness, cortico-amygdalar networks, and externalizing behaviors in healthy children. Biol Psychiatry 75, 65–72. https://doi.org/10.1016/j.biopsych.2013.06.008

Avants, B. B., Libon, D. J., Rascovsky, K., Boller, A., McMillan, C. T., Massimo, L., …& Grossman, M. (2014). Sparse canonical correlation analysis relates network-level atrophy to multivariate cognitive measures in a neurodegenerative population. Neuroimage, 84, 698–711. https://doi.org/10.1016/j.neuroimage.2013.09.048

Biederman, J., DiSalvo, M., Vaudreuil, C., Wozniak, J., Uchida, M., Yvonne Woodworth, K., Green, A., Faraone, S. V., 2020. Can the Child Behavior Checklist (CBCL) help characterize the types of psychopathologic conditions driving child psychiatry referrals? Scand J Child Adolesc Psychiatr Psychol 8, 57–165. https://doi.org/10.21307/sjcapp-2020-016

Boekel, W., Wagenmakers, E.J., Belay, L., Verhagen, J., Brown, S. and Forstmann, B.U., 2015. A purely confirmatory replication study of structural brain-behavior correlations. Cortex, 66, pp.115–133. https://doi.org/10.1016/j.cortex.2014.11.019

Burgaleta, M., Johnson, W., Waber, D.P., Colom, R., Karama, S., 2014. Cognitive ability changes and dynamics of cortical thickness development in healthy children and adolescents. Neuroimage 84, 810–819. https://doi.org/10.1016/j.neuroimage.2013.09.038

Button, K.S., Ioannidis, J.P.A., Mokrysz, C., Nosek, B.A., Flint, J., Robinson, E.S.J., Munafò, M.R., 2013. Power failure: Why small sample size undermines the reliability of neuroscience. Nat Rev Neurosci 14, 365–376. https://doi.org/10.1038/nrn3475

Byrd, A.L., Hawes, S.W., Waller, R., Delgado, M.R., Sutherland, M.T., Dick, A.S., Trucco, E.M., Riedel, M.C., Pacheco-Colón, I., Laird, A.R. and Gonzalez, R., 2021. Neural response to monetary loss among youth with disruptive behavior disorders and callous-unemotional traits in the ABCD study. NeuroImage: Clinical, 32, p.102810. https://doi.org/10.1016/j.nicl.2021.102810

Casey, B.J., Cannonier, T., Conley, M.I., Cohen, A.O., Barch, D.M., Heitzeg, M.M., Soules, M.E., Teslovich, T., Dellarco, D. v., Garavan, H., Orr, C.A., Wager, T.D., Banich, M.T., Speer, N.K., Sutherland, M.T., Riedel, M.C., Dick, A.S., Bjork, J.M., Thomas, K.M., Chaarani, B., Mejia, M.H., Hagler, D.J., Daniela Cornejo, M., Sicat, C.S., Harms, M.P., Dosenbach, N.U.F., Rosenberg, M., Earl, E., Bartsch, H., Watts, R., Polimeni, J.R., Kuperman, J.M., Fair, D.A., Dale, A.M., 2018. The Adolescent Brain Cognitive Development (ABCD) study: Imaging acquisition across 21 sites. Dev Cogn Neurosci 32, 43–54. https://doi.org/10.1016/j.dcn.2018.03.001

Churchill, N., Spring, R., Abdi, H., Kovacevic, N., Mcintosh, A.R., Strother, S., n.d. 2013. The Stability of Behavioral PLS Results in Ill-Posed Neuroimaging Problems.

Dale, A.M., Fischl, B., Sereno, M.I., 1999. Cortical surface-based analysis. I. Segmentation and surface reconstruction. Neuroimage 9, 179–194.

Desikan, R.S., Ségonne, F., Fischl, B., Quinn, B.T., Dickerson, B.C., Blacker, D., Buckner, R.L., Dale, A.M., Maguire, R.P., Hyman, B.T., Albert, M.S., Killiany, R.J., 2006. An automated labeling system for subdividing the human cerebral cortex on MRI scans into gyral based regions of interest. Neuroimage 31, 968–980. https://doi.org/10.1016/j.neuroimage.2006.01.021

de Winter, J. C. F., Gosling, S. D., & Potter, J. (2016). Comparing the Pearson and Spearman correlation coefficients across distributions and sample sizes: A tutorial using simulations and empirical data. Psychological Methods, 21(3), 273–290. https://doi.org/10.1037/met0000079

Drysdale, A.T., Grosenick, L., Downar, J., Dunlop, K., Mansouri, F., Meng, Y., Fetcho, R.N., Zebley, B., Oathes, D.J., Etkin, A., Schatzberg, A.F., Sudheimer, K., Keller, J., Mayberg, H.S., Gunning, F.M., Alexopoulos, G.S., Fox, M.D., Pascual-Leone, A., Voss, H.U., Casey, B.J., Dubin, M.J., Liston, C., 2017. Resting-state connectivity biomarkers define neurophysiological subtypes of depression. Nat Med 23, 28–38. https://doi.org/10.1038/nm.4246

Ducharme, S., Albaugh, M.D., Hudziak, J.J., Botteron, K.N., Nguyen, T.V., Truong, C., Evans, A.C., Karama, S., 2014. Anxious/depressed symptoms are linked to right ventromedial prefrontal cortical thickness maturation in healthy children and young adults. Cerebral Cortex 24, 2941– 2950. https://doi.org/10.1093/cercor/bht151

Ducharme, S., Hudziak, J.J., Botteron, K.N., Ganjavi, H., Lepage, C., Collins, D.L., Albaugh, M.D., Evans, A.C., Karama, S., 2011. Right anterior cingulate cortical thickness and bilateral striatal volume correlate with child behavior checklist aggressive behavior scores in healthy children. Biol Psychiatry 70, 283–290. https://doi.org/10.1016/j.biopsych.2011.03.015

Ehrlich, S., Brauns, S., Yendiki, A., Ho, B.C., Calhoun, V., Charles Schulz, S., Gollub, R.L., Sponheim, S.R., 2012. Associations of cortical thickness and cognition in patients with schizophrenia and healthy controls. Schizophr Bull 38, 1050–1062. https://doi.org/10.1093/schbul/sbr018

Fischl, B., Dale, A.M., 2000. Measuring the thickness of the human cerebral cortex from magnetic resonance images. Proc Natl Acad Sci U S A 97, 11050–11055.

Grady, C.L., Rieck, J.R., Nichol, D., Rodrigue, K.M., Kennedy, K.M., 2021. Influence of sample size and analytic approach on stability and interpretation of brain-behavior correlations in task-related fMRI data. Hum Brain Mapp 42, 204–219. https://doi.org/10.1002/hbm.25217

Hagler, D.J., Hatton, S.N., Makowski, C., Cornejo, M.D., Fair, D.A., Dick, A.S., Sutherland, M.T., Casey, B.J., Barch, D.M., Harms, M.P., Watts, R., Bjork, J.M., Garavan, H.P., Hilmer, L., Pung, C.J., Sicat, C.S., Kuperman, J., Bartsch, H., Xue, F., Heitzeg, M.M., Laird, A.R., Trinh, T.T., Gonzalez, R., Tapert, S.F., Riedel, M.C., Squeglia, L.M., Hyde, L.W., Rosenberg, M.D., Earl, E.A., Howlett, K.D., Baker, F.C., Soules, M., Diaz, J., de Leon, O.R., Thompson, W.K., Neale, M.C., Herting, M., Sowell, E.R., Alvarez, R.P., Hawes, S.W., Sanchez, M., Bodurka, J., Breslin, F.J., Morris, A.S., Paulus, M.P., Simmons, W.K., Polimeni, J.R., der Kouwe, A. van, Nencka, A.S., Gray, K.M., Pierpaoli, C., Matochik, J.A., Noronha, A., Aklin, W.M., Conway, K., Glantz, M., Hoffman, E., Little, R., Lopez, M., Pariyadath, V., Weiss, S.R.B., Wolff-Hughes, D.L., DelCarmen-Wiggins, R., Ewing, S.W.F., Miranda-Dominguez, O., Nagel, B.J., Perrone, A.J., Sturgeon, D.T., Goldstone, A., Pfefferbaum, A., Pohl, K.M., Prouty, D., Uban, K., Bookheimer, S.Y., Dapretto, M., Galvan, A., Bagot, K., Giedd, J., Infante, M.A., Jacobus, J., Patrick, K., Shilling, P.D., Desikan, R., Li, Y., Sugrue, L., Banich, M.T., Friedman, N., Hewitt, J.K., Hopfer, C., Sakai, J., Tanabe, J., Cottler, L.B., Nixon, S.J., Chang, L., Cloak, C., Ernst, T., Reeves, G., Kennedy, D.N., Heeringa, S., Peltier, S., Schulenberg, J., Sripada, C., Zucker, R.A., Iacono, W.G., Luciana, M., Calabro, F.J., Clark, D.B., Lewis, D.A., Luna, B., Schirda, C., Brima, T., Foxe, J.J., Freedman, E.G., Mruzek, D.W., Mason, M.J., Huber, R., McGlade, E., Prescot, A., Renshaw, P.F., Yurgelun-Todd, D.A., Allgaier, N.A., Dumas, J.A., Ivanova, M., Potter, A., Florsheim, P., Larson, C., Lisdahl, K., Charness, M.E., Fuemmeler, B., Hettema, J.M., Steinberg, J., Anokhin, A.P., Glaser, P., Heath, A.C., Madden, P.A., Baskin-Sommers, A., Constable, R.T., Grant, S.J., Dowling, G.J., Brown, S.A., Jernigan, T.L., Dale, A.M., 2018. Image processing and analysis methods for the Adolescent Brain Cognitive Development Study. bioRxiv. https://doi.org/10.1101/457739

Helmer, M., Warrington, S., Mohammadi-Nejad, A. R., Ji, J. L., Howell, A., Rosand, B., … & Murray, J. D. (2020). On stability of canonical correlation analysis and partial least squares with application to brain-behavior associations. BioRxiv, 2020-08. https://doi.org/10.1101/2020.08.25.265546

Hill, W.D., Hagenaars, S.P., Marioni, R.E., Harris, S.E., Liewald, D.C.M., Davies, G., Okbay, A., McIntosh, A.M., Gale, C.R., Deary, I.J., 2016. Molecular Genetic Contributions to Social Deprivation and Household Income in UK Biobank. Current Biology 26, 3083–3089. https://doi.org/10.1016/j.cub.2016.09.035

Hotelling, H., Relations between two sets of variates. Biometrika, 1936. 28(3-4): p. 321–377

Huang-Pollock, C., Shapiro, Z., Galloway-Long, H., Weigard, A., 2017. Is Poor Working Memory a Transdiagnostic Risk Factor for Psychopathology? J Abnorm Child Psychol 45, 1477–1490. https://doi.org/10.1007/s10802-016-0219-8

Huber, P. J. (2011). Robust statistics. In International encyclopedia of statistical science (pp. 1248–1251). Springer, Berlin, Heidelberg.

Ing, A., Sämann, P.G., Chu, C., Tay, N., Biondo, F., Robert, G., Jia, T., Wolfers, T., Desrivières, S., Banaschewski, T., Bokde, A.L.W., Bromberg, U., Büchel, C., Conrod, P., Fadai, T., Flor, H., Frouin, V., Garavan, H., Spechler, P.A., Gowland, P., Grimmer, Y., Heinz, A., Ittermann, B., Kappel, V., Martinot, J.L., Meyer-Lindenberg, A., Millenet, S., Nees, F., van Noort, B., Orfanos, D.P., Martinot, M.L.P., Penttilä, J., Poustka, L., Quinlan, E.B., Smolka, M.N., Stringaris, A., Struve, M., Veer, I.M., Walter, H., Whelan, R., Andreassen, O.A., Agartz, I., Lemaitre, H., Barker, E.D., Ashburner, J., Binder, E., Buitelaar, J., Marquand, A., Robbins, T.W., Schumann, G., 2019. Identification of neurobehavioural symptom groups based on shared brain mechanisms. Nat Hum Behav 3, 1306–1318. https://doi.org/10.1038/s41562-019-0738-8

Itahashi, T., Fujino, J., Sato, T., Ohta, H., Nakamura, M., Kato, N., Hashimoto, R.-I., di Martino, A., Aoki, Y.Y., 2020. Neural correlates of shared sensory symptoms in autism and attention-deficit/hyperactivity disorder. Brain Commun 2. https://doi.org/10.1093/braincomms/fcaa186

Jacobs, G., Voineskos, A., Hawco, C., Stefanik, L., Forde, N., Dickie, E., Lai, M.-C., Szatmari, P., Schachar, R., Crosbie, J., Arnold, P., Goldenberg, A., Erdman, L., Lerch, J., Anagnostou, E., Ameis, S., 2020. Integration of Brain and Behavior Measures for Identification of Data-Driven Groups Cutting Across Children with ASD, ADHD, or OCD. https://doi.org/10.1101/2020.02.11.944744

Jacobson, L. A., Murphy-Bowman, S. C., Pritchard, A. E., Tart-Zelvin, A., Zabel, T. A., & Mahone, E. M. (2012). Factor structure of a sluggish cognitive tempo scale in clinically-referred children. Journal of abnormal child psychology, 40, 1327–1337. https://doi.org/10.1007/s10802-012-9643-6

Kebets, V., Holmes, A.J., Orban, C., Tang, S., Li, J., Sun, N., Kong, R., Poldrack, R.A., Yeo, B.T.T., 2019. Somatosensory-Motor Dysconnectivity Spans Multiple Transdiagnostic Dimensions of Psychopathology. Biol Psychiatry 86, 779–791. https://doi.org/10.1016/j.biopsych.2019.06.013

S. Kotz, N. Johnson (Eds.), Encyclopedia of Statistical Sciences, Wiley, New York (1985), pp. 581-591

Krishnan, A., Williams, L.J., McIntosh, A.R., Abdi, H., 2011. Partial Least Squares (PLS) methods for neuroimaging: A tutorial and review. Neuroimage 56, 455–475. https://doi.org/10.1016/j.neuroimage.2010.07.034

Lawson, G.M., Duda, J.T., Avants, B.B., Wu, J., Farah, M.J., 2013. Associations between children’s socioeconomic status and prefrontal cortical thickness. Dev Sci 16, 641–652. https://doi.org/10.1111/desc.12096

Lombardo, M. v., Lai, M.C., Baron-Cohen, S., 2019. Big data approaches to decomposing heterogeneity across the autism spectrum. Mol Psychiatry 24, 1435–1450. https://doi.org/10.1038/s41380-018-0321-0

Mardia, K., J. Kent, and J. Bibby, Multivariate Analysis. 1979, New York: Academic Press.

Mahony, B.W., Tu, D., Rau, S., Liu, S., Lalonde, F.M., Alexander-Bloch, A.F., Satterthwaite, T.D., Shinohara, R.T., Bassett, D.S., Milham, M.P., Raznahan, A., 2022. IQ Modulates Coupling Between Diverse Dimensions of Psychopathology in Children and Adolescents. J Am Acad Child Adolesc Psychiatry. https://doi.org/10.1016/j.jaac.2022.06.015

Marek, A.S., Tervo-clemmens, B., Calabro, F.J., David, F., Uriarte, J., Snider, K., Tam, A., Chen, J., Dillan, J., Greene, D.J., Petersen, S.E., Nichols, T.E., Thomas, B.T., 2020. Towards Reproducible Brain-Wide Association Studies.

Marek, S., Tervo-Clemmens, B., Nielsen, A.N., Wheelock, M.D., Miller, R.L., Laumann, T.O., Earl, E., Foran, W.W., Cordova, M., Doyle, O. and Perrone, A., 2019. Identifying reproducible individual differences in childhood functional brain networks: An ABCD study. Developmental cognitive neuroscience, 40, p.100706. https://doi.org/10.1016/j.dcn.2019.100706

Masouleh, S.K., Eickhoff, S.B., Hoffstaedter, F., Genon, S. and Alzheimer’s Disease Neuroimaging Initiative, 2019. Empirical examination of the replicability of associations between brain structure and psychological variables. elife, 8, p.e43464. https://doi.org/10.7554/eLife.43464

McIntosh, A.R., 2021. Comparison of Canonical Correlation and Partial Least Squares analyses of simulated and empirical data. https://doi.org/10.48550/arXiv.2107.06867

McIntosh, A.R., Lobaugh, N.J., 2004. Partial least squares analysis of neuroimaging data: Applications and advances. Neuroimage 23, 250–263. https://doi.org/10.1016/j.neuroimage.2004.07.020

Mihalik, A., Ferreira, F.S., Rosa, M.J., Moutoussis, M., Ziegler, G., Monteiro, J.M., Portugal, L., Adams, R.A., Romero-Garcia, R., Vértes, P.E., Kitzbichler, M.G., Váša, F., Vaghi, M.M., Bullmore, E.T., Fonagy, P., Goodyer, I.M., Jones, P.B., Hauser, T., Neufeld, S., Clair, M.S., Whitaker, K., Inkster, B., Prabhu, G., Ooi, C., Toseeb, U., Widmer, B., Bhatti, J., Villis, L., Alrumaithi, A., Birt, S., Bowler, A., Cleridou, K., Dadabhoy, H., Davies, E., Firkins, A., Granville, S., Harding, E., Hopkins, A., Isaacs, D., King, J., Kokorikou, D., Maurice, C., McIntosh, C., Memarzia, J., Mills, H., O’Donnell, C., Pantaleone, S., Scott, J., Fearon, P., Suckling, J., van Harmelen, A.L., Kievit, R., Dolan, R., Mourão-Miranda, J., 2019. Brain-behaviour modes of covariation in healthy and clinically depressed young people. Sci Rep 9. https://doi.org/10.1038/s41598-019-47277-3

Modabbernia, A., Janiri, D., Doucet, G.E., Reichenberg, A., Frangou, S., 2021. Multivariate Patterns of Brain-Behavior-Environment Associations in the Adolescent Brain and Cognitive Development Study. Biol Psychiatry 89, 510–520. https://doi.org/10.1016/j.biopsych.2020.08.014

Moser, D.A., Doucet, G.E., Lee, W.H., Rasgon, A., Krinsky, H., Leibu, E., Ing, A., Schumann, G., Rasgon, N., Frangou, S., 2018. Multivariate associations among behavioral, clinical, and multimodal imaging phenotypes in patients with psychosis. JAMA Psychiatry 75, 386–395. https://doi.org/10.1001/jamapsychiatry.2017.4741

Myers, L. and Sirois, M.J., 2006. Spearman correlation coefficients, differences between. Encyclopedia of statistical sciences, 12. https://doi.org/10.1002/0471667196.ess5050.pub2

Nakua, H., Hawco, C., Forde, N. J., Jacobs, G. R., Joseph, M., Voineskos, A. N., … & Ameis, S. H. (2022). Cortico-amygdalar connectivity and externalizing/internalizing behavior in children with neurodevelopmental disorders. Brain structure and function, 227(6), 1963–1979. https://doi.org/10.1007/s00429-022-02483-0

Nikolaidis, A., Chen, A.A., He, X., Shinohara, R., Vogelstein, J., Milham, M. and Shou, H., 2022. Suboptimal phenotypic reliability impedes reproducible human neuroscience. bioRxiv. https://doi.org/10.1101/2022.07.22.501193

Nour, M.M., Liu, Y., Dolan, R.J., 2022. Functional neuroimaging in psychiatry and the case for failing better. Neuron 110, 2524–2544. https://doi.org/10.1016/j.neuron.2022.07.005

Ooi, L.Q.R., Chen, J., Zhang, S., Kong, R., Tam, A., Li, J., Dhamala, E., Zhou, J.H., Holmes, A.J., Yeo, B.T.T., 2022. Comparison of individualized behavioral predictions across anatomical, diffusion and functional connectivity MRI. Neuroimage 263, 119636. https://doi.org/10.1016/j.neuroimage.2022.119636

Orlhac, F., Eertink, J.J., Cottereau, A.S., Zijlstra, J.M., Thieblemont, C., Meignan, M., Boellaard, R. and Buvat, I., 2022. A guide to ComBat harmonization of imaging biomarkers in multicenter studies. Journal of Nuclear Medicine, 63,.172–179. https://doi.org/10.2967/jnumed.121.262464

Owens, M.M., Potter, A., Hyatt, C.S., Albaugh, M., Thompson, W.K., Jernigan, T., Yuan, D., Hahn, S., Allgaier, N. and Garavan, H., 2021. Recalibrating expectations about effect size: A multi-method survey of effect sizes in the ABCD study. PloS one 16, p.e0257535. https://doi.org/10.1371/journal.pone.0257535

Piccolo, L.R., Merz, E.C., He, X., Sowell, E.R., Noble, K.G., 2016. Age-related differences in cortical thickness vary by socioeconomic status. PLoS One 11, 1–18. https://doi.org/10.1371/journal.pone.0162511

Poldrack, R.A., Baker, C.I., Durnez, J., Gorgolewski, K.J., Matthews, P.M., Munafò, M.R., Nichols, T.E., Poline, J.B., Vul, E., Yarkoni, T., 2017. Scanning the horizon: Towards transparent and reproducible neuroimaging research. Nat Rev Neurosci 18, 115–126. https://doi.org/10.1038/nrn.2016.167

Qin, S., Young, C.B., Duan, X., Chen, T., Supekar, K., Menon, V., 2014. Amygdala subregional structure and intrinsic functional connectivity predicts individual differences in anxiety during early childhood. Biol Psychiatry 75, 892–900. https://doi.org/10.1016/j.biopsych.2013.10.006

Rakesh, D., Zalesky, A., Whittle, S., 2021. Similar but distinct – Effects of different socioeconomic indicators on resting state functional connectivity: Findings from the Adolescent Brain Cognitive Development (ABCD) Study®. Dev Cogn Neurosci 51, 101005. https://doi.org/10.1016/j.dcn.2021.101005

Romer, A.L., Elliott, M.L., Knodt, A.R., Sison, M.L., Ireland, D., Houts, R., Ramrakha, S., Poulton, R., Keenan, R., Melzer, T.R., Moffitt, T.E., Caspi, A., Hariri, A.R., 2021. Pervasively thinner neocortex as a transdiagnostic feature of general psychopathology. American Journal of Psychiatry 178, 174–182. https://doi.org/10.1176/appi.ajp.2020.19090934

Ronan, L., Alexander-Bloch, A., Fletcher, P.C., 2020. Childhood Obesity, Cortical Structure, and Executive Function in Healthy Children. Cerebral Cortex 30, 2519–2528. https://doi.org/10.1093/cercor/bhz257

Scheinost, D., Noble, S., Horien, C., Greene, A.S., Lake, E.M., Salehi, M., Gao, S., Shen, X., O’Connor, D., Barron, D.S., Yip, S.W., Rosenberg, M.D., Constable, R.T., 2019. Ten simple rules for predictive modeling of individual differences in neuroimaging. Neuroimage 193, 35–45. https://doi.org/10.1016/j.neuroimage.2019.02.057

Seok, D., Beer, J., Jaskir, M., Smyk, N., Jaganjac, A., Makhoul, W., Cook, P., Elliott, M., Shinohara, R., Sheline, Y.I., 2021. Differential impact of transdiagnostic, dimensional psychopathology on multiple scales of functional connectivity. https://doi.org/10.1101/2021.03.05.434151

Smith, S.M., Nichols, T.E., Vidaurre, D., Winkler, A.M., Behrens, T.E.J., Glasser, M.F., Ugurbil, K., Barch, D.M., Van Essen, D.C., Miller, K.L., 2015. A positive-negative mode of population covariation links brain connectivity, demographics and behavior. Nat Neurosci. https://doi.org/10.1038/nn.4125

Storch, E. A., Murphy, T. K., Bagner, D. M., Johns, N. B., Baumeister, A. L., Goodman, W. K., & Geffken, G. R. (2006). Reliability and validity of the child behavior checklist obsessive-compulsive scale. Journal of Anxiety Disorders, 20(4), 473–485. https://doi.org/10.1016/j.janxdis.2005.06.002

Stoycos, S.A., Piero, L. Del, Margolin, G., Kaplan, J.T., Saxbe, D.E., 2017. Neural correlates of inhibitory spillover in adolescence: Associations with internalizing symptoms. Soc Cogn Affect Neurosci 12, 1637–1646. https://doi.org/10.1093/scan/nsx098

Thompson, W. K., Barch, D. M., Bjork, J. M., Gonzalez, R., Nagel, B. J., Nixon, S. J., & Luciana, M. (2019). The structure of cognition in 9 and 10 year-old children and associations with problem behaviors: Findings from the ABCD study’s baseline neurocognitive battery. Developmental cognitive neuroscience, 36, 100606. https://doi.org/10.1016/j.dcn.2018.12.004

Vijayakumar, N., Allen, N.B., Dennison, M., Byrne, M.L., Simmons, J.G., Whittle, S., 2017. Cortico-amygdalar maturational coupling is associated with depressive symptom trajectories during adolescence. Neuroimage 156, 403–411. https://doi.org/10.1016/j.neuroimage.2017.05.051

Walter, M., Alizadeh, S., Jamalabadi, H., Lueken, U., Dannlowski, U., Walter, H., Olbrich, S., Colic, L., Kambeitz, J., Koutsouleris, N., Hahn, T., Dwyer, D.B., 2019. Translational machine learning for psychiatric neuroimaging. Prog Neuropsychopharmacol Biol Psychiatry 91, 113–121. https://doi.org/10.1016/j.pnpbp.2018.09.014

Wang, H. T., Bzdok, D., Margulies, D., Craddock, C., Milham, M., Jefferies, E., & Smallwood, J. (2018). Patterns of thought: Population variation in the associations between large-scale network organisation and self-reported experiences at rest. Neuroimage, 176, 518–527. https://doi.org/10.1016/j.neuroimage.2018.04.064

Weintraub, S., Dikmen, S. S., Heaton, R. K., Tulsky, D. S., Zelazo, P. D., Bauer, P. J., … & Gershon, R. C. (2013). Cognition assessment using the NIH Toolbox. Neurology, 80(11 Supplement 3), S54–S64. https://doi.org/10.1212/WNL.0b013e3182872ded

Wilde, E.A., Merkley, T.L., Bigler, E.D., Max, J.E., Schmidt, A.T., Ayoub, K.W., McCauley, S.R., Hunter, J. V., Hanten, G., Li, X., Chu, Z.D., Levin, H.S., 2012. Longitudinal changes in cortical thickness in children after traumatic brain injury and their relation to behavioral regulation and emotional control. International Journal of Developmental Neuroscience 30, 267–276. https://doi.org/10.1016/j.ijdevneu.2012.01.003

Winter, J.C.F. De, Gosling, S.D., Potter, J., 2016. Supplemental Material for Comparing the Pearson and Spearman Correlation Coefficients Across Distributions and Sample Sizes: A Tutorial Using Simulations and Empirical Data. Psychol Methods 21, 273–290. https://doi.org/10.1037/met0000079.supp

Witten, D. M., & Tibshirani, R. (2009). Covariance-regularized regression and classification for high dimensional problems. Journal of the Royal Statistical Society: Series B (Statistical Methodology), 71(3), 615–636. https://doi.org/10.1111/j.1467-9868.2009.00699.x

Xia, C.H., Ma, Z., Ciric, R., Gu, S., Betzel, R.F., Kaczkurkin, A.N., Calkins, M.E., Cook, P.A., García de la Garza, A., Vandekar, S.N., Cui, Z., Moore, T.M., Roalf, D.R., Ruparel, K., Wolf, D.H., Davatzikos, C., Gur, R.C., Gur, R.E., Shinohara, R.T., Bassett, D.S., Satterthwaite, T.D., 2018. Linked dimensions of psychopathology and connectivity in functional brain networks. Nat Commun 9, 1–14. https://doi.org/10.1038/s41467-018-05317-y

Yip, S.W., Kiluk, B., Scheinost, D., 2020. Toward Addiction Prediction: An Overview of Cross-Validated Predictive Modeling Findings and Considerations for Future Neuroimaging Research. Biol Psychiatry Cogn Neurosci Neuroimaging 5, 748–758. https://doi.org/10.1016/j.bpsc.2019.11.001

Ziegler, G., Dahnke, R., Winkler, A.D., Gaser, C., 2013. Partial least squares correlation of multivariate cognitive abilities and local brain structure in children and adolescents. Neuroimage 82, 284–294. https://doi.org/10.1016/j.neuroimage.2013.05.088

