## Supplemental Materials for "Comparing the stability and reproducibility of brain-behaviour relationships found using Canonical Correlation Analysis and Partial Least Squares within the ABCD Sample"

### 1. Explanation of Canonical Correlations Analysis (CCA) and Partial Least Squares Correlations (PLS)

Both CCA and PLS approaches aim to address a maximization problem: searching for largest linear relationships between two data matrices (e.g., **X** and **Y**) based on specific constraints. The maximization problem is solved using the singular value decomposition (SVD) which is a technique that decomposes and extracts underlying dimensions (called *latent variables*) from a rectangular matrix. In PLS and CCA, the SVD decomposes the following matrices:

**PLS:**

$$\mathbf{R}_{\mathbf{XY}} = \mathbf{X}^{\mathsf{T}}\mathbf{Y} \text{ (i.e., the cross-product matrix),}$$

**CCA:**

$$\mathbf{\Omega} = (\mathbf{X}^{\mathsf{T}}\mathbf{X})^{-0.5}\,\mathbf{X}^{\mathsf{T}}\mathbf{Y}(\mathbf{Y}^{\mathsf{T}}\mathbf{Y})^{-0.5}.$$

The SVD solves the following maximization problem for each latent variable (denoted by $\ell$):

**PLS:**

$$s_\ell = \arg\max_{\mathbf{u}_\ell,\mathbf{v}_\ell} \{\mathbf{u}_\ell^{\mathsf{T}}\mathbf{R}_{\mathbf{XY}}\mathbf{v}_\ell\} \quad \text{such that} \quad \mathbf{u}_\ell^{\mathsf{T}}\mathbf{u}_\ell = \mathbf{v}_\ell^{\mathsf{T}}\mathbf{v}_\ell = 1.$$

**CCA:**

$$s_\ell = \arg\max_{\mathbf{u}_\ell,\mathbf{v}_\ell} \{\mathbf{u}_\ell^{\mathsf{T}}\mathbf{\Omega}\mathbf{v}_\ell\} \quad \text{such that} \quad \mathbf{u}_\ell^{\mathsf{T}}\mathbf{u}_\ell = \mathbf{v}_\ell^{\mathsf{T}}\mathbf{v}_\ell = 1.$$

*Note:* **u** *and* **v** *are the left and the right singular vectors, and* **s** *are the singular value of the decomposed matrix (where* $s_1 \geq s_2 \geq \cdots \geq s_\ell \geq \cdots \geq s_L$*). The superscript* $^{\mathsf{T}}$ *indicates the transpose of a matrix or a vector such that rows become columns.*

The maximization problem in CCA and PLS can be solved iteratively by a gradient descent algorithm such that the first pair of singular vectors identified ($\mathbf{u}_1$ and $\mathbf{v}_1$) explain the maximum amount of variance in $\mathbf{R}_{\mathbf{XY}}$ or $\mathbf{\Omega}$. In this way, the $\mathbf{u}_1$ and $\mathbf{v}_1$ singular vectors store the coefficients of each variable (analogous to beta weights in a linear regression), respectively of **X** and **Y**, that form the first latent variable. These coefficients are called *loadings*. The singular value $s_1$ gives the standard deviation, therefore quantifies the variance, of the first latent variable. The second pair of singular vectors ($\mathbf{u}_2$ and $\mathbf{v}_2$) and singular value ($s_2$) are identified to be orthogonal to the first pair (i.e., $\mathbf{u}_1^{\mathsf{T}}\mathbf{u}_2 = \mathbf{v}_1^{\mathsf{T}}\mathbf{v}_2 = 0$) and explain the maximum amount of the remaining variance from the first latent variable with $s_1 \geq s_2$. This process continues until all variance of $\mathbf{R_{XY}}$ or $\mathbf{\Omega}$ is explained by all latent variables. With $s_1 \geq s_2 \geq \ldots \geq s_\ell \geq \ldots \geq s_L$, the SVD ensures that the relationship between the original **X** and **Y** matrices is explained by the latent variables in a descending order with the strongest relationship (explains the greatest amount of variance) in the first latent variable.

In CCA, the singular vectors (**U** and **V**) from the SVD are further reweighted by the Cholesky decomposition of the respective within-block correlation matrix. This obtains *beta weights* which are analogous to beta weights in a linear regression. As a result, the singular values used in CCA are generalized singular values. Throughout the paper, we have referred to them as singular vectors to remain consistent with the language used for PLS. The mathematical derivation of the beta weights are explained below:

$$\mathbf{SVD(\Omega) = USV^T}$$
$$\mathbf{U_{beta} = (X^TX)^{-0.5}\, U}$$
$$\mathbf{V_{beta} = (Y^TY)^{-0.5}\, V}$$

For the convenience of comparing CCA and PLS, we will refer to $\mathbf{U_{beta}}$ and $\mathbf{V_{beta}}$ of CCA from here on and in the main paper as **U** and **V**, respectively.

Formally, this decomposition can be expressed in one step as:

**PLS:**

$$\mathbf{R_{XY} = USV^T} \quad \text{such that} \quad \mathbf{U^TU = V^TV = I}$$

**CCA:**

$$\mathbf{\Omega = USV^T} \quad \text{such that} \quad \mathbf{U^T(X^TX)U = V^T(Y^TY)V = I},$$

Where the **U** and **V** are the matrices of the left and the right singular vectors, **S** is a diagonal matrix with $s_\ell$ on the diagonal and 0s on the off diagonal, and **I** denotes the identity matrix that has 1s on the diagonal and 0s on the off diagonal. The left singular vector matrix **U** has *n* left singular vectors (**u)** on the columns and describes the loadings of **X** for all latent variables. The right singular vector matrix **V** has *n* right singular vectors (**v)** on the columns and describes the loadings of the **Y** matrix for all latent variables. The singular values consist of the effect sizes of the multivariate relationship; in PLS, the *covariance* and, in CCA, the *correlation*. In the CCA literature, this correlation is also referred to as the canonical correlation (often denoted by $\delta$).

The critical difference between CCA and PLS is the maximization of correlation versus covariance. This maximization occurs to the relationship between the latent scores $\mathbf{L_X}$ and $\mathbf{L_Y}$. These latent scores are projections of the matrices **X** and **Y** onto the latent dimensions by multiplying their original variables by their respective singular vectors **U** and **V**. These scores are mathematically expressed as:

**PLS:**

$$\mathbf{L_X} = \mathbf{XU}$$

$$\mathbf{L_Y} = \mathbf{YV}$$

$$\text{where } \mathbf{S} = \mathbf{L_X}^T\mathbf{L_Y} = \mathbf{U}^T(\mathbf{X}^T\mathbf{Y})\mathbf{V};$$

$$\text{for each latent variable, } s_\ell = \boldsymbol{l}_X^T\boldsymbol{l}_Y$$

**CCA:**

$$\mathbf{L_X} = \mathbf{XU}$$

$$\mathbf{L_Y} = \mathbf{YV}$$

$$\text{where } \mathbf{S} = (\mathbf{X}^T\mathbf{X})^{-0.5}\,\mathbf{L_X}^T\mathbf{L_Y}(\mathbf{Y}^T\mathbf{Y})^{-0.5};$$

$$\text{for each latent variable, } s_\ell = \frac{l_X^T l_Y^T}{\sqrt{X^T X} \times \sqrt{Y^T Y}}$$

*Note: In PLS and CCA, the $\mathbf{L}_X$ and $\mathbf{L}_Y$ matrix is the product of a direct multiplication of the original scores (**X/Y**) and loadings (**U/V**). In PLS, the singular values in **S** are maximized by obtaining the cross-product between $\mathbf{L}_X$ and $\mathbf{L}_Y$ (i.e., covariance). In CCA, $\mathbf{L}_X$ and $\mathbf{L}_Y$ are computed the same way as in PLS; however, the singular values of the cross-product (i.e.,$\mathbf{L}_X^T\mathbf{L}_Y$) are pre- and post-multiplied by the inverse of the square-root of the within-block correlations of **X** and **Y** (i.e., $(\mathbf{X}^T\mathbf{X})^{-\frac{1}{2}}$ and $(\mathbf{Y}^T\mathbf{Y})^{-\frac{1}{2}}$). This equation of computing **S** in CCA is equivalent to computing the correlation between $\mathbf{L}_X$ and $\mathbf{L}_Y$.*

### *1.2. Structure Coefficients for CCA*

In addition to the beta weights, McIntosh et al. (2020) implemented CCA by further defining and analyzing *structure coefficients*. The structure coefficients are calculated by multiplying the within-block correlation matrices of **X** and **Y** by their respective beta weights. This step reintroduces the variance of **X** and **Y** so that the LVs generated using these structural coefficients are more similar to the LVs from PLS. The mathematical derivation of the structural coefficients are explained below:

$$\mathbf{U}_{\mathbf{structCoef}} = (\mathbf{X}^{\mathbf{T}}\mathbf{X})(\mathbf{X}^{\mathbf{T}}\mathbf{X})^{-0.5}\ \mathbf{U}$$

$$\mathbf{V}_{\mathbf{structCoef}} = (\mathbf{Y}^{\mathbf{T}}\mathbf{Y})(\mathbf{Y}^{\mathbf{T}}\mathbf{Y})^{-0.5}\ \mathbf{V}$$

*Note: structCoef = structure coefficients*

For the main paper, we interpreted the LVs generated from the beta weights across all analyses. We included the CCA results for the first analysis with the structural coefficients in *Figure S4*.

### 2. Analytical Decisions

#### *2.1. Spearman versus Pearson correlation matrix*

Although the main paper presented the CCA and PLS results when decomposing a Spearman cross-correlation matrix, Pearson correlations are more commonly implemented. As such, we also explored whether using a specific correlation coefficient would alter the overall results of the first analysis (relationship between cortical thickness and CBCL scores). We found that the identified LVs for both CCA and PLS were similar when implementing a Spearman versus a Pearson cross-correlation matrix (see *Figure 2; Figure S2*), suggesting a consistent LV for linear and monotonic relationships between **X** and **Y**. When decomposing a Pearson correlation matrix, there were 6 and 7 statistically significant LVs for the CCA and PLS models, respectively. Further, performing the split-half resampling for the PLS model using a Pearson cross-correlation matrix revealed cortical thickness loadings for $LV_1$ that just surpassed the reproducibility threshold ($z$-score = 2.04). This suggests that the relationships found when using a Pearson cross-product matrix may be stronger compared to using a Spearman's cross-product matrix. However, given the possibility of non-linear relationships between the cortical thickness and CBCL data, the Pearson cross-correlation may inflate the correlations. Prior work showed that using a Spearman's correlation coupled with a Fisher's $z$-transform yielded more robust results compared to using Pearson correlation coefficients on non-normal data (Myers & Sirios, 2006). As such, the relationships assessed using a Spearman's correlation are likely more accurate compared to Pearson's correlation in this sample.

#### *2.2. Transforming the Behavioural Data*

Given the skewness of the CBCL data, we attempted to transform the data to impose normality of the distribution. We used the log transform, as used in prior reports (Dienes et al. 2002; Gross et al. 2006; Tollenaar et al. 2021), however, it did not remove the skewness of the CBCL data. As a result, we decided to use the Spearman's correlation matrix (without transforming the data) to address the skewness of the CBCL scores.

#### *2.3. Arbitrary Sign Flip Correction for Bootstrap Resampling*

In the bootstrap resampling process, there is a possibility of reflections (i.e., sign flips) in the resampled matrix each time the SVD is performed. These sign flips should be corrected to reduce the estimation bias of the bootstrap resampling (McIntosh & Lobaugh, 2004). In the bootstrap resampling analysis, we generated 1000 singular vector matrices, and assessed whether the signs of the elements in each singular vector were arbitrarily flipped. This was done by multiplying each generated **U** or **V** matrix by the respective empirical **U** or **V** matrix and determining whether the diagonals of the product matrix (the product of $\mathbf{U}_{\mathbf{generated}}$ and $\mathbf{U}_{\mathbf{empirical}}$ or $\mathbf{V}_{\mathbf{generated}}$ and $\mathbf{V}_{\mathbf{empirical}}$) were negative. If the value on the diagonal was negative, then that singular vector would be multiplied by -1 to correct the sign flip. From there, we calculated the 95% confidence interval of each variable in $LV_1$ from the 1000 generated **U** and **V** matrices.

### 3. Sensitivity Analyses

#### *3.1. Subset of sample without head injuries*

To obtain this subsample, we removed participants with a "yes" (coded as 1) for the following variables: *medhx_6i* (head injury), *medhx_2c* (seizure), *medhx_2m* (multiple sclerosis), *medhx_2h* (epilepsy), *medhx_2f* (cerebral palsy), *medhx_2c* (brain injury) from the *abcd_mx01.csv* as part of the tabulated data from ABCD. Overall, for both the first and second analyses, the brain-behaviour relationship identified in $LV_1$ when using the main sample is preserved in this subsample for both CCA and PLS suggesting that this relationship is robust against possible effects of history of brain injury (see *Figure S3*; *Figure S4*)).

#### *3.2. Subset of sample regressed for household income data*

$LV_1$ for both CCA and PLS models in this subsample are consistent in the first and second analytical sample (see *Figure S3*; *Figure S4*). This suggests that the brain-behaviour relationship identified in $LV_1$ for CCA and PLS is robust against variation in SES.

#### *3.3. Including head size as a regressor*

To reduce some of the multicollinearity among the cortical thickness measures, we included total brain volume as a covariate in the linear regression performed prior to conducting the CCA and PLS analyses. The use of regressors when examining cortical thickness is not consistent among prior work; some have covaried for whole brain volumetric measures in addition to age and sex (Zhu et al. 2021; Ameis et al. 2016; Hall et al. 2021), others only covary for age and sex (Owens et al. 2021), and some covary for age-squared, age x sex, age-squared x sex (Zhu et al. 2021; Modabbernia et al. 2021). Given the limited age range of the current study, we decided not to include age-squared in the models given that there was a high collinearity between age-squared and age. Further, the relationship between age and cortical thickness or psychopathology was best described by a linear model (see *Table S8* for examples). As such, for the main analyses, we added total cortical volume as a regression in the linear regression model prior to performing the CCA or PLS. To ensure that covarying for total cortical volume was not substantially influencing the results, we compared the cross-product matrices ($\mathbf{R_{XY}}$ and $\mathbf{\Omega}$) when total cortical volume was covaried for and when it was not (but age, sex, site, and scanner were covaried for). When correlating these two matrices, we found high correlations ($r > .8$) suggesting that regressing out total cortical volume has limited influence on the subsequent analyses. See *Figure S8* for the correlation plot which depicts the Pearson correlations between $\mathbf{R_{XY}}$ and $\mathbf{\Omega}$ from the main sample when total cortical volume is and is not regressed. The diagonal of the correlation plot indicates that the majority of variables within the brain and behavioural matrix are very similar whether total cortical volume is regressed or not.

### 4. Exploratory analyses

#### *4.1. Subsample with Higher Psychopathology*

In the subset of the sample with an elevated CBCL total score (*t*-score > 60; $n = 1016$), the cross-block correlations in the $\mathbf{R_{XY}}$ and $\mathbf{\Omega}$ matrices were larger than that of the main sample (*Figure S5*). The singular values were also higher than the main sample ($LV_1$: PLS = .67, CCA = .33). None of the LVs were statistically significant or reproducible across the CCA and PLS analysis.

#### *4.2. Subsample with endorsement of all CBCL subscales*

Our second post-hoc analysis examined a subset of the ABCD participants who had some endorsement of each CBCL subscale score (i.e., no value of 0; $n = 5196$). This analysis was performed to assess whether the 0-inflation of the CBCL scores was driving the low within-method generalizability of the results. The results revealed similar brain-behaviour relationships as the main sample (singular values of $LV_1$: PLS = .3, CCA = .15; *Figure S6*). None of the LVs were statistically significant or reproducible.

### 5. Figures

*Figure S1. Consort Diagram describing the number and reason of participant exclusion from the analysis.*

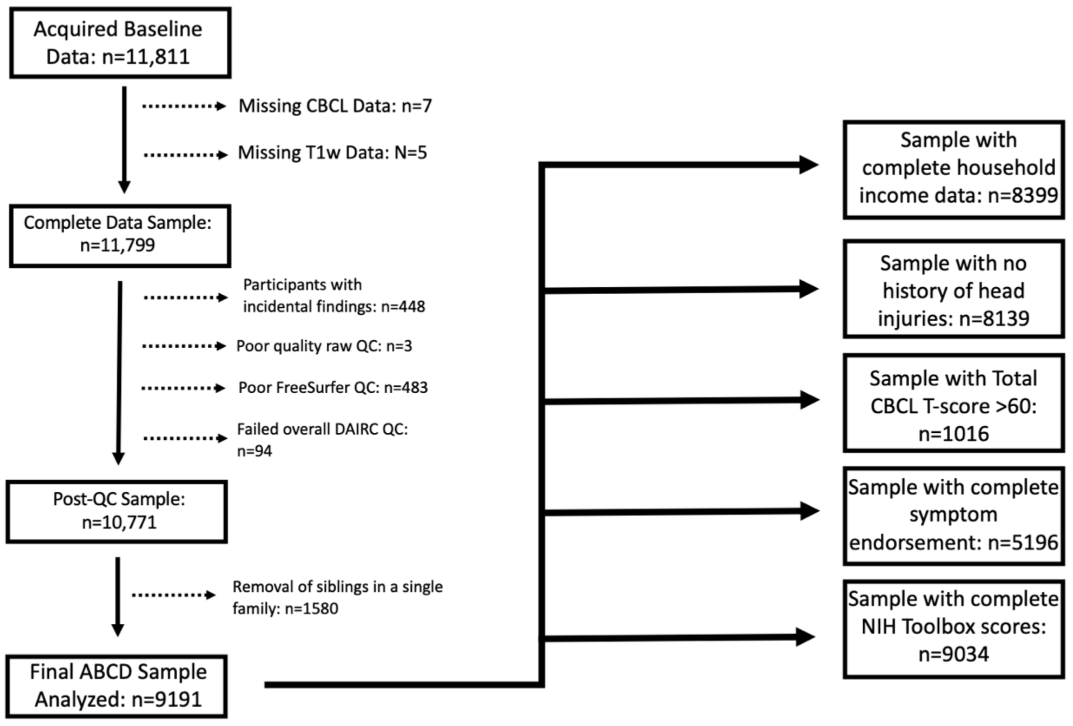

*Note: The variables used to exclude participants are as follows. The incidental findings variable was mrif_score and participants were excluded if they were considered to "need clinical referral" or "immediate clinical referral". Participants with a zero value for the iqc_t1_ok_ser variable were excluded indicating that they had poor quality raw T1-weighted scans. Participants were excluded if they received a "reject" from the fsqc_qc variable indicating they failed FreeSurfer QC. Finally, any additional participants who failed the DAIRC QC were excluded (received a zero for the imgincl_t1w_include variable). The ABCD dataset includes data collected from siblings, twins and triplets as part of the sample. To reduce multicollinearity, we only retained one sibling per family.*

*Figure S2. CCA and PLS results when implementing a Pearson cross-correlation matrix to examine the relationship between cortical thickness and CBCL scores.*

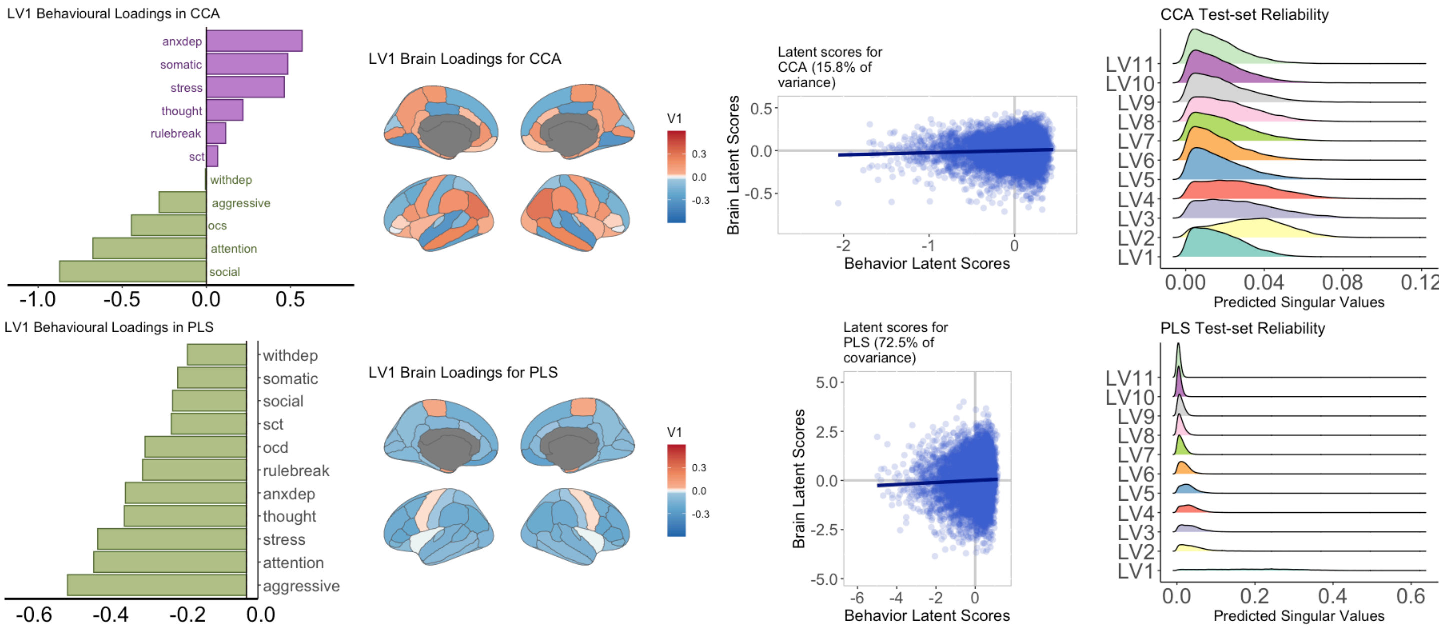

Note: Unthresholded behaviour and brain loadings from the PLS and CCA analysis performed in the main sample using a Pearson cross-correlation matrix for **Rxy** and **Ω**. Overall, the relationships in $LV_1$ using a Pearson cross-correlation matrix are similar to those using a Spearman cross-correlation matrix. The LVs were not stable for CCA and PLS. Prior to calculating the latent scores, the brain and behavioural loadings have been standardized by the singular values. OCD = obsessive compulsive disorder (symptoms), withdep = withdrawn/depression symptoms, sct = sluggish-cognitive-tempo, anxdep = anxiety/depression symptoms, rulebreak = rule breaking behaviour.

*Figure S3. CCA and PLS loadings for the two sensitivity analyses; controlling for household income (SES Subset) and removing participants without head injuries (No Head Injury Subset) in the first analysis (between CBCL and cortical thickness)*

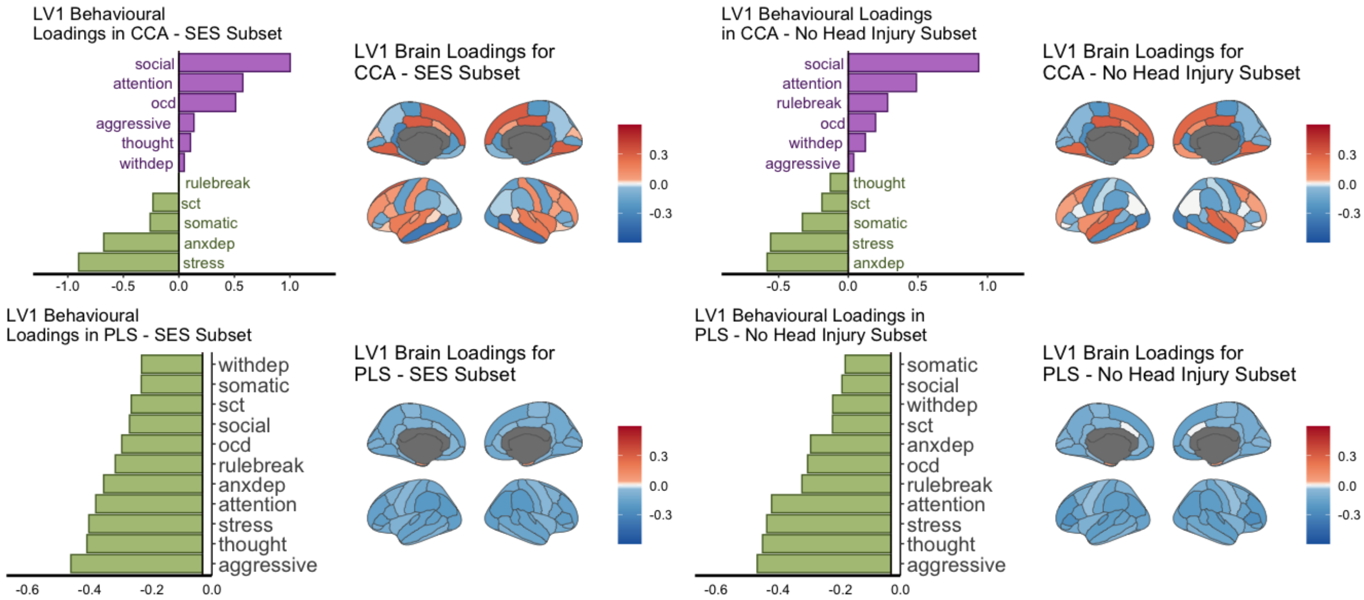

Note: In both sensitivity analysis subsets, we find similar relationships to the first analysis from the main manuscript. Social problems and aggressive behaviours are the highest contributing behavioural variable in $LV_1$ for CCA and PLS, respectively. We find overall covariation between behavioural and brain loadings in CCA (i.e., positive and negative loadings in the behavioural measures linked to positive and negative loadings in the brain measures). In the PLS analysis, we find the same trend such that decreased behavioural loadings is linked to decreased brain loadings (i.e., lower behavioural problems are linked to decreased cortical thickness). OCD = obsessive compulsive disorder (symptoms), withdep = withdrawn/depression symptoms, sct = sluggish-cognitive-tempo, anxdep = anxiety/depression symptoms, rulebreak = rule breaking behaviour.

*Figure S4. $LV_1$ CCA loadings for structure coefficients in the first analysis (between CBCL and cortical thickness)*

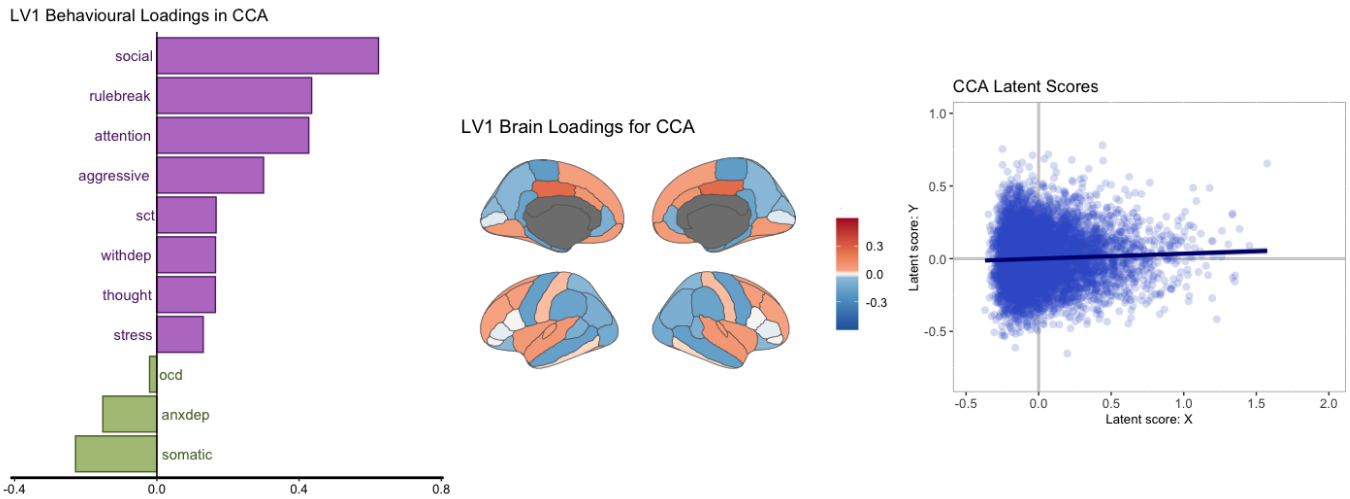

Note: A spearman cross-correlation matrix was used in this analysis. The pattern of $LV_1$ is similar to that when using the generalized singular vectors from the SVD (called beta weights). Social problems remain the highest behavioural loading and we find patterns of covariation between the behaviour and brain loadings. Prior to calculating the latent scores, the brain and behavioural loadings have been standardized by the singular values. OCD = obsessive compulsive disorder (symptoms), withdep = withdrawn/depression symptoms, sct = sluggish-cognitive-tempo, anxdep = anxiety/depression symptoms, rulebreak = rule breaking behaviour. $LV_1$ when using the structural coefficients accounted for 19.3% of the variance.

*Figure S5. $LV_1$ CCA and PLS loadings for the elevated-CBCL subsample (CBCL t-score > 60, n = 1016).*

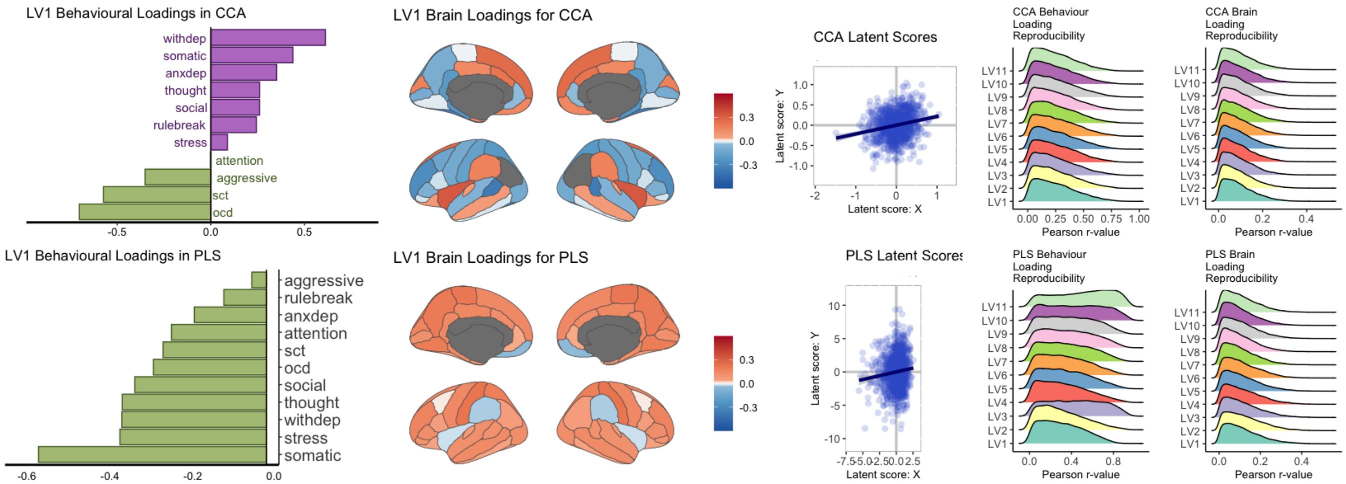

Note: Unthresholded behaviour and brain weights from the PLS and CCA analysis performed in the subsample with elevated CBCL scores. Overall, the brain-behaviour relationships in $LV_1$ differed between the CCA and PLS analyses in the main sample. Although the CCA analysis depicts the covariation trend, withdrawn/depression symptoms have the highest behavioural loading. The PLS analysis depicts a similar homogeneous relationship, however, less behavioural problems (i.e., lower CBCL scores) is linked to increased cortical thickness. Prior to calculating the latent scores, the brain and behavioural loadings have been standardized by the singular values. $LV_1$ for CCA accounted for 14.7% of the variance, and $LV_1$ for PLS accounted for 53.9% of the covariance. OCD = obsessive compulsive disorder (symptoms), withdep = withdrawn/depression symptoms, sct = sluggish-cognitive-tempo, anxdep = anxiety/depression symptoms, rulebreak = rule breaking behaviour.

*Figure S6. $LV_1$ CCA and PLS loadings for the subsample with full subscale symptom endorsement (n = 5196)*

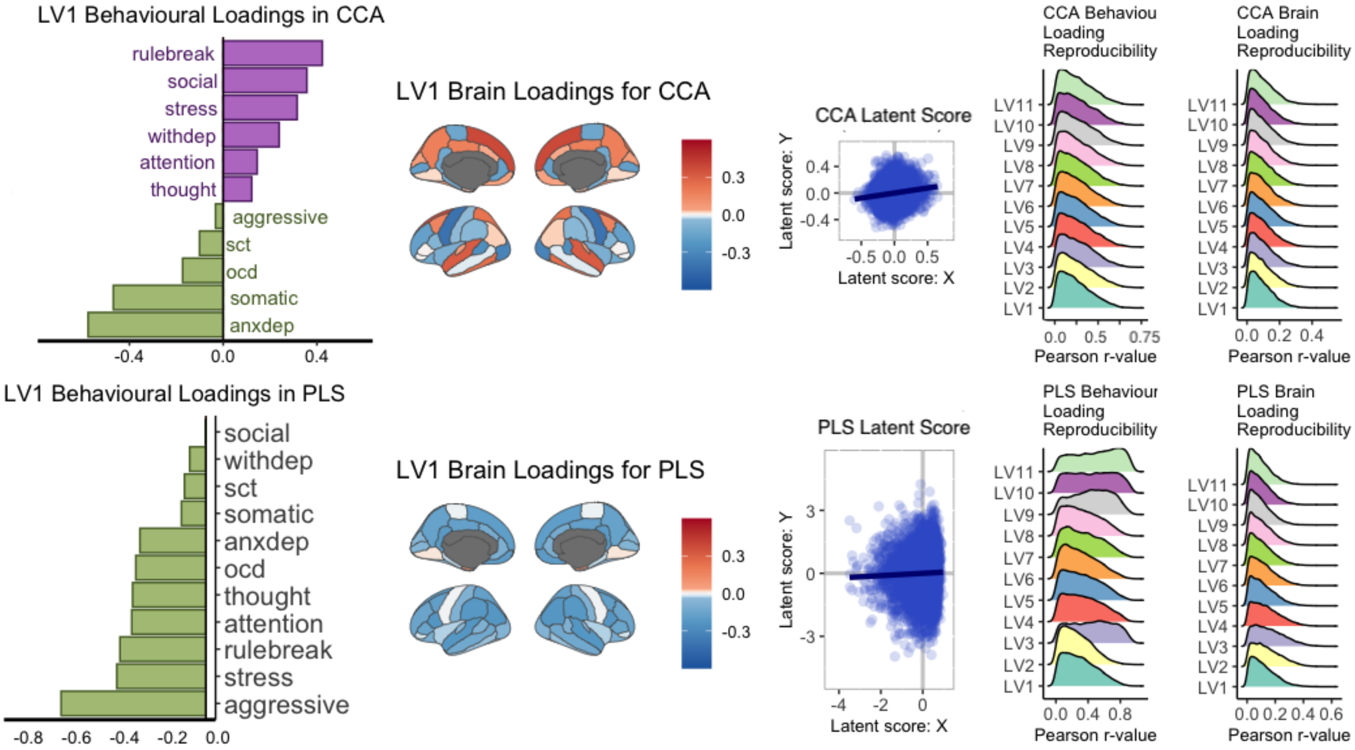

Note: Unthresholded behaviour and brain loadings from the PLS and CCA analysis performed in the subsample with subscale endorsement for each of the CBCL subscales. Overall, the brain-behaviour relationships found in $LV_1$ in this subsample are similar to that found in the primary analysis. Prior to calculating the latent scores, the brain and behavioural loadings have been standardized by the singular values. $LV_1$ for CCA accounted for 16% of the variance, and LV1 for PLS accounted for 52.3% of the covariance. OCD = obsessive compulsive disorder (symptoms), withdep = withdrawn/depression symptoms, sct = sluggish-cognitive-tempo, anxdep = anxiety/depression symptoms, rulebreak = rule breaking behaviour.

*Figure S7. CCA and PLS analytical results for the two sensitivity analyses in the second analysis (between cortical thickness and cognitive performance); controlling for household income (SES Subset) and removing participants without head injuries (No Head Injury Subset).*

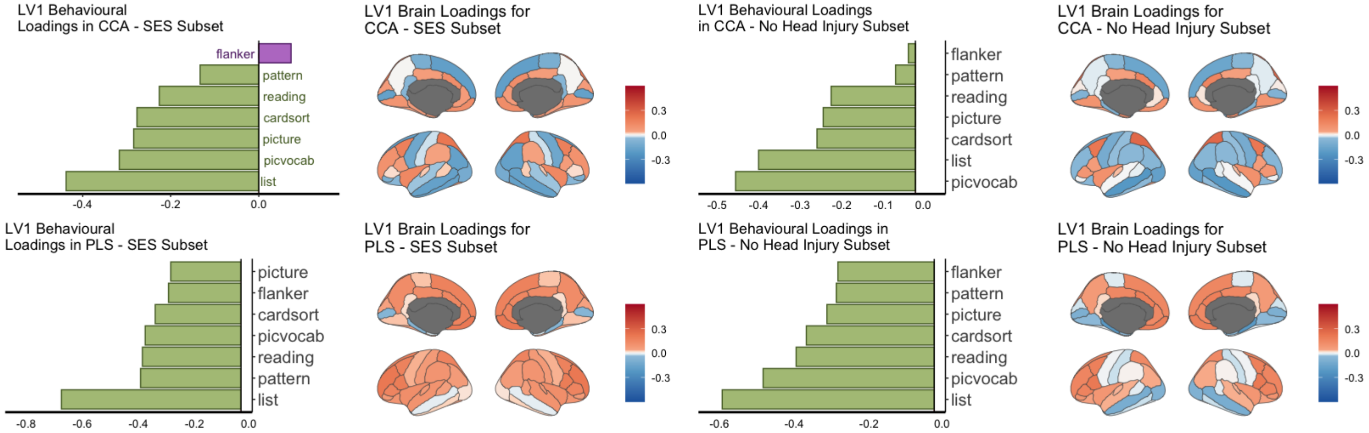

Note: In both sensitivity analysis subsets, we find overall similar relationships to the results of the second analysis included in the main manuscript. One difference is the positive loading of the Flanker task in the CCA when regressing out household income. Performance on the list sorting working memory task is consistently the top contributing variable from the NIH scores (except in the CCA when removing participants with no head injuries). We find overall covariation in the brain loadings in CCA and PLS (i.e., positive and negative loadings), however PLS results show more positive loadings. Flanker = Flanker Task, pattern = pattern comparison processing speed task, cardsort = dimensional change card sort task, reading = oral reading recognition task, picture = picture vocabulary task, list = list sorting working memory task, picvocab = picture vocabulary task.

*Figure S8. Correlation plot depicting correlations between the cross-product matrices (*$\mathbf{R_{XY}}$ *and* $\boldsymbol{\Omega}$*) that include total cortical volume as a regressor and that do not.*

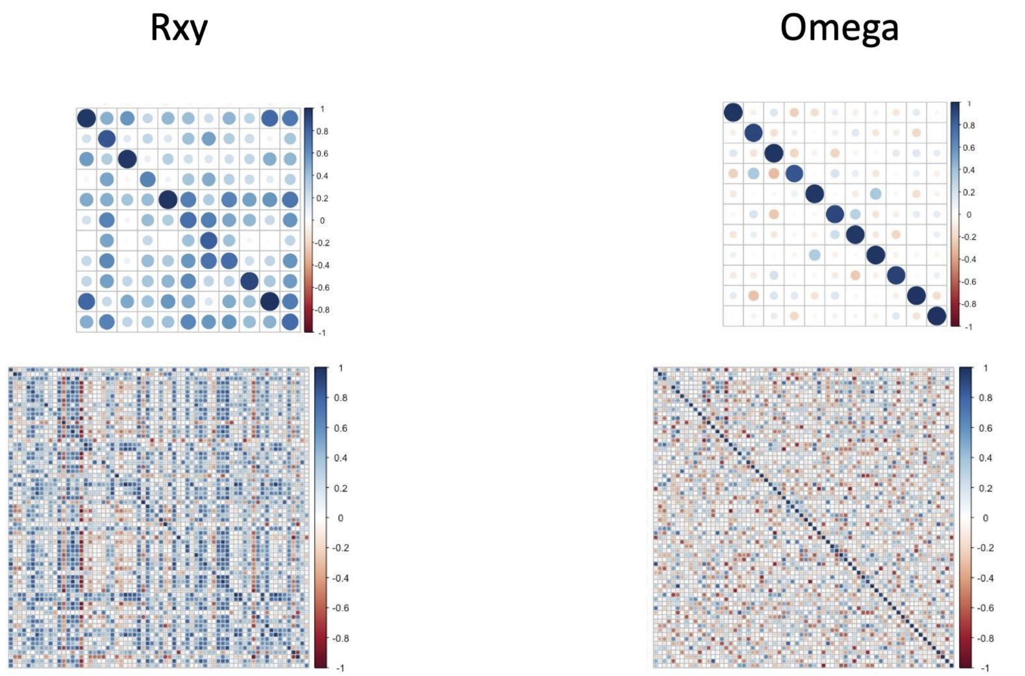

Note: The diagonal of the $\mathbf{R_{XY}}$ and $\mathbf{\Omega}$ matrices across the different brain and behavioural measures are all $r > .8$ suggesting that the relationships identified would be similar regardless of regressing out total brain volume or not.

*Figure S9. Within- and between-block correlations of the first analysis (examining CCA and PLS analysis between CBCL subscale scores and cortical thickness).*

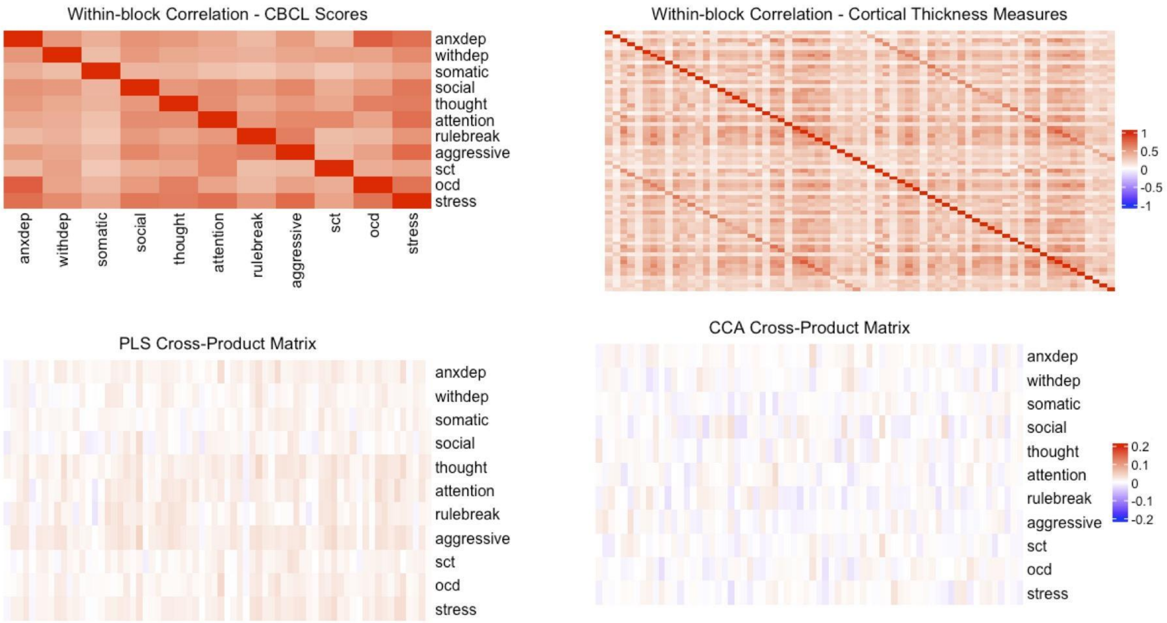

Note: OCD = obsessive compulsive disorder (symptoms), withdep = withdrawn/depression symptoms, sct = sluggish-cognitive-tempo, anxdep = anxiety/depression symptoms, rulebreak = rule breaking behaviour. PLS cross-product matrix = $\mathbf{R_{XY}}$, CCA cross-product matrix = $\mathbf{\Omega}$.

*Figure S10. Within- and between-block correlations of the second analysis (examining CCA and PLS analysis between NIH Cognitive Toolbox scores and cortical thickness).*

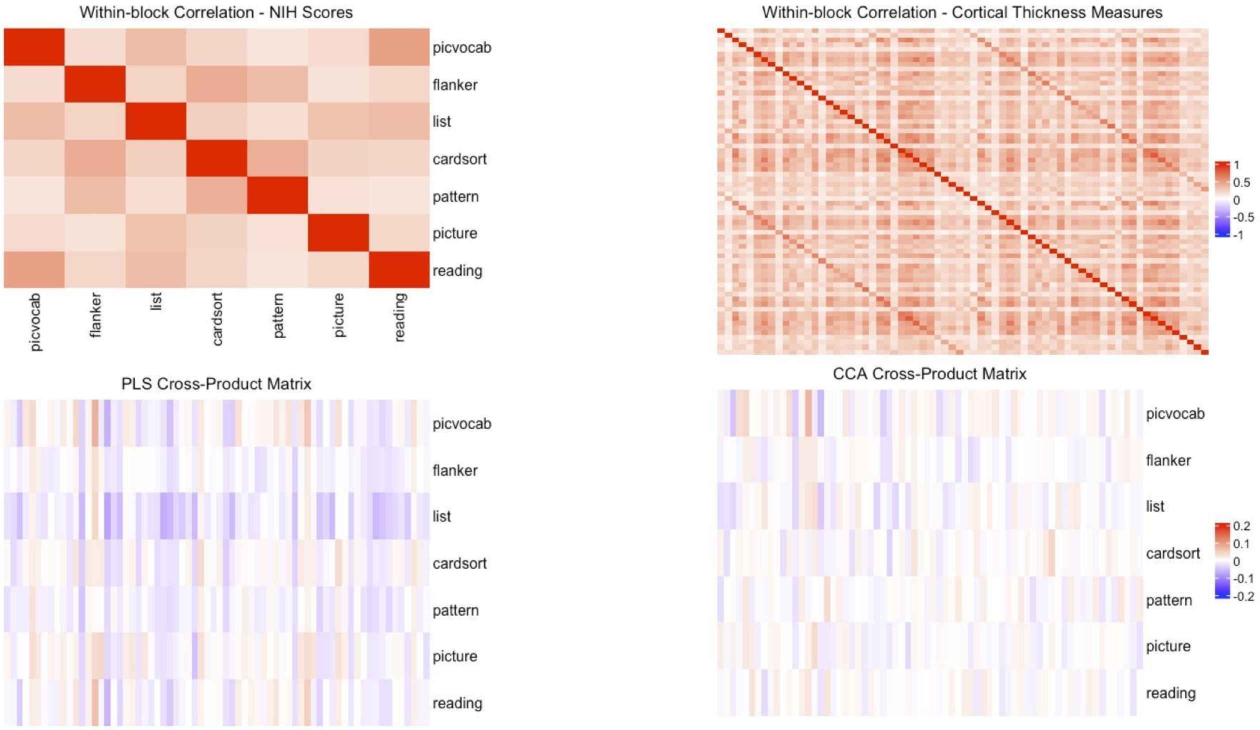

Note: Flanker = Flanker Task, pattern = pattern comparison processing speed task, cardsort = dimensional change card sort task, reading = oral reading recognition task, picture = picture vocabulary task, list = list sorting working memory task, picvocab = picture vocabulary task. PLS cross-product matrix = $\mathbf{R_{XY}}$, CCA cross-product matrix = $\mathbf{\Omega}$.

*Figure S11. Within- and between-block correlations of the post-hoc subsample with elevated CBCL scores (CBCL Total T-score >60; n=1016).*

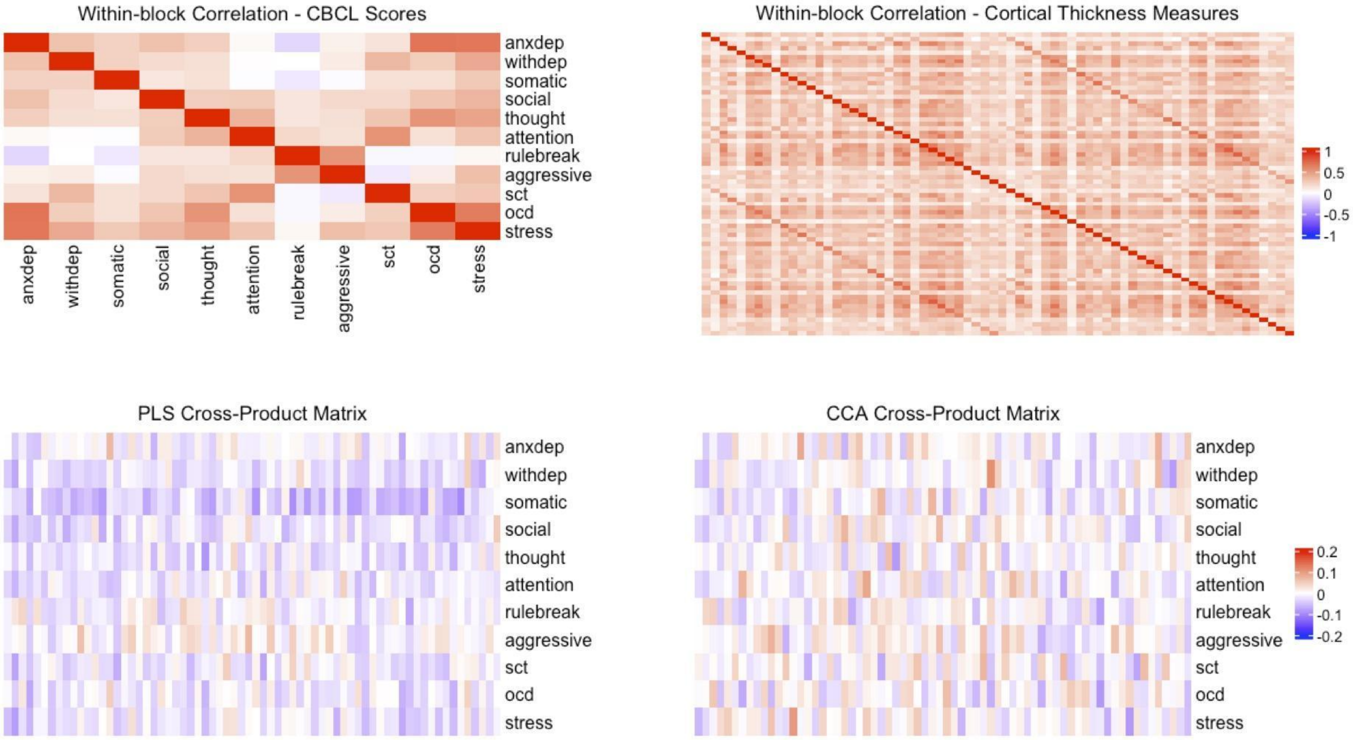

Note: OCD = obsessive compulsive disorder (symptoms), withdep = withdrawn/depression symptoms, sct = sluggish-cognitive-tempo, anxdep = anxiety/depression symptoms, rulebreak = rule breaking behaviour. PLS cross-product matrix = **Rxy**, CCA cross-product matrix = **Ω**.

*Figure S12. Distributions of residuals of the behavioural variables used in the first, second, and post-hoc analyses.*

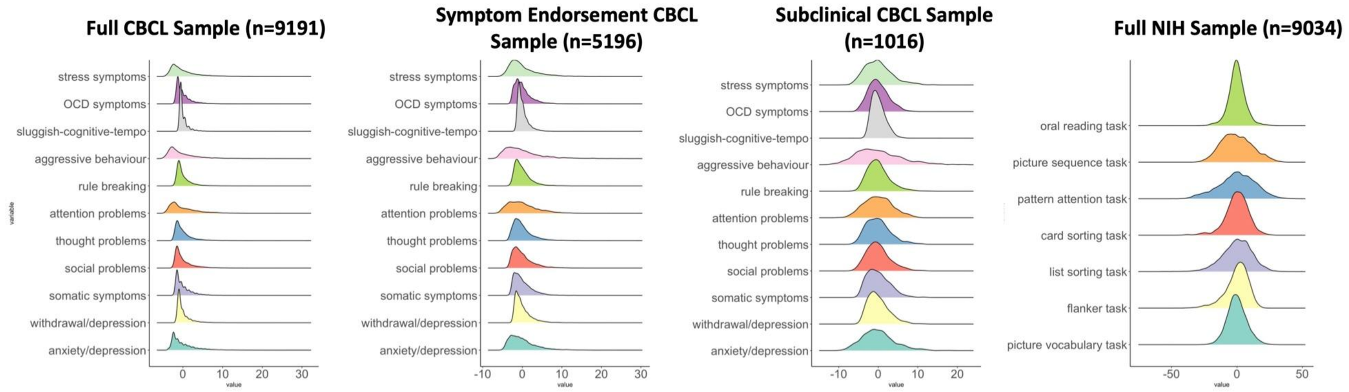

*Figure S13. Barplots depicting the loadings of the CBCL and cortical thickness elements in $LV_1$ of the first analysis.*

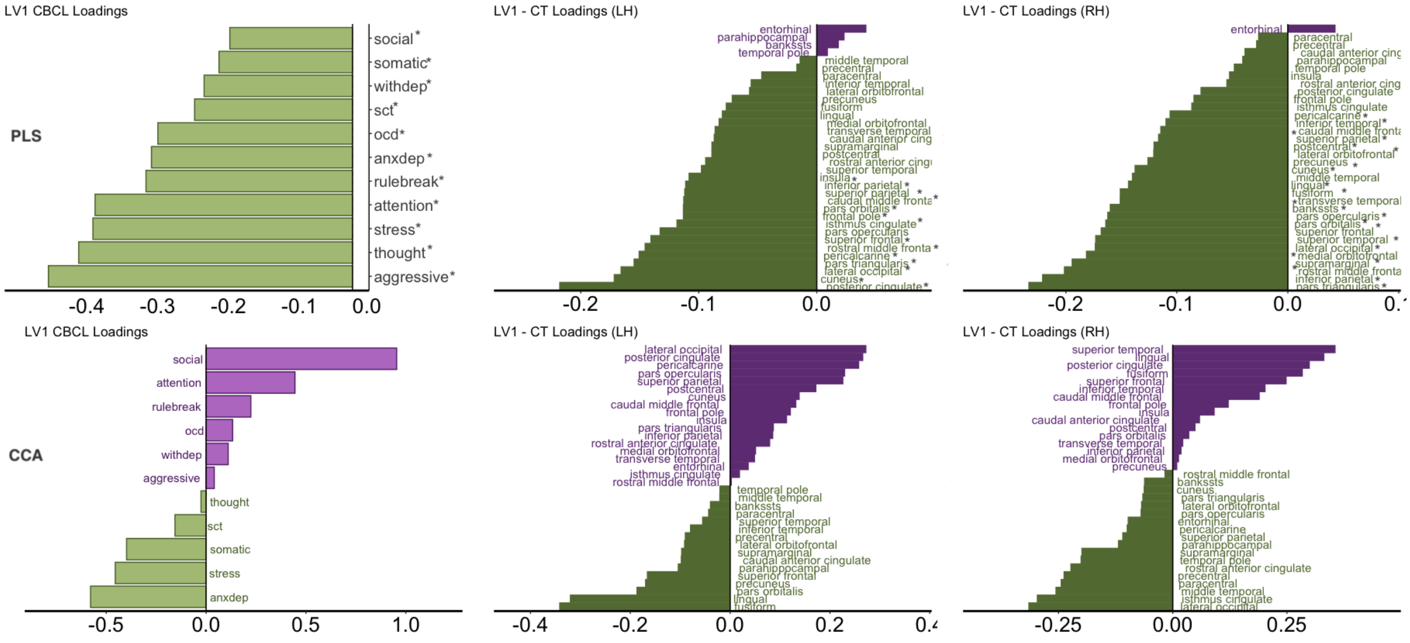

Note: Asterisks indicate that the element was stable as assessed by bootstrap resampling (i.e., the 95% confidence interval did not include zero). The asterisk may be after or before the name of the element. OCD = obsessive compulsive disorder (symptoms), withdep = withdrawn/depression symptoms, sct = sluggish-cognitive-tempo, anxdep = anxiety/depression symptoms, rulebreak = rule breaking behaviour.

*Figure S14. Barplots depicting the loadings of the NIH and cortical thickness elements in $LV_1$ of the second analysis.*

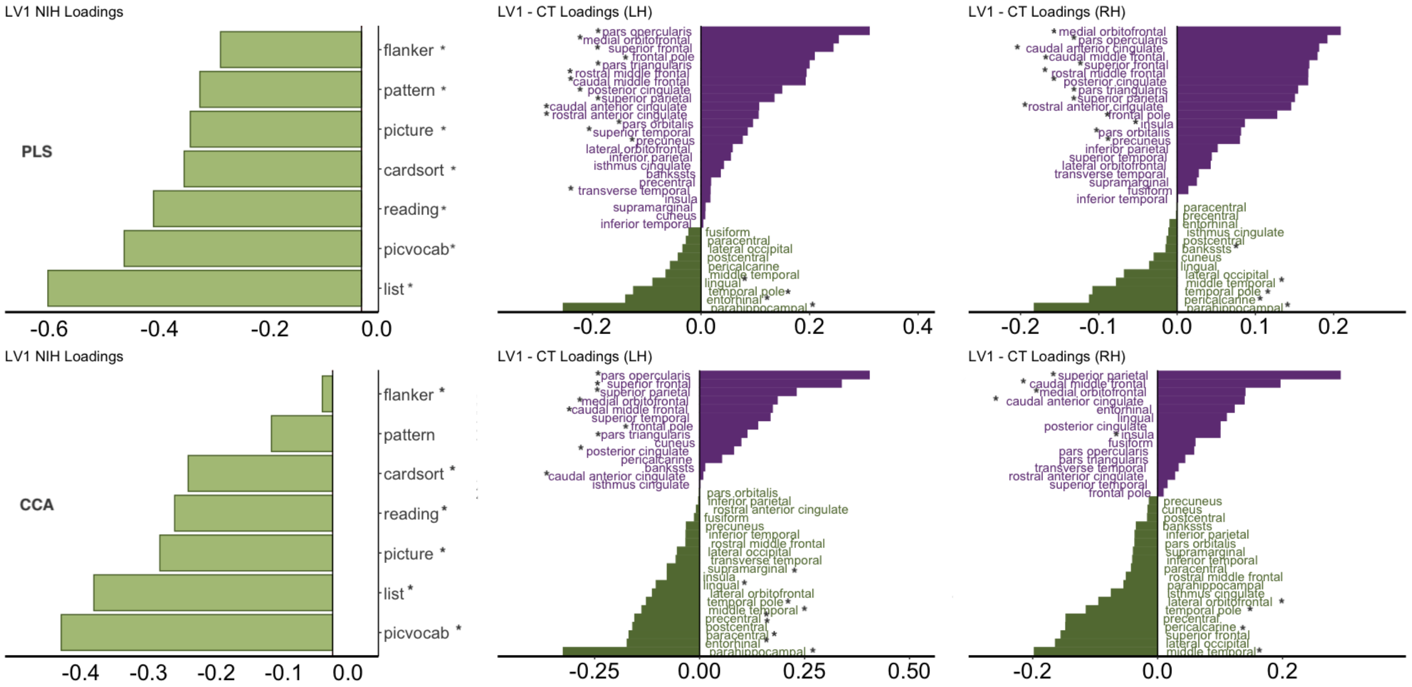

Note: Asterisks indicate that the element was stable as assessed by bootstrap resampling (i.e., the 95% confidence interval did not cross zero). The asterisk may be after or before the name of the element. Flanker = Flanker Task, pattern = pattern comparison processing speed task, cardsort = dimensional change card sort task, reading = oral reading recognition task, picture = picture vocabulary task, list = list sorting working memory task, picvocab = picture vocabulary task.

### 6. Tables

*Table S1. Demographic Characteristics of the ABCD main sample analyzed and the SES and head injury subsamples*

| | **1. Included ABCD Sample (n=9191)** | | **2. SES Sample (n=8399)** | | **3. No Head Injury Sample (n=8139)** | | **F-score, p-value** |
|---|---|---|---|---|---|---|---|
| | **Mean [range]** | **SD** | **Mean [range]** | **SD** | **Mean [range]** | **SD** | |
| Age (in months) | 118.9 [107-133] | 7.4 | 118.9 [107-133] | 7.4 | 118.9 [107-133] | 7.4 | F=0.041, p=0.96 |
| CBCL Total Score (raw) | 18.56 [0-139] | 18.22 | 18.49 [0-139] | 18.16 | 17.7 [0-139] | 17.4 | F=5.38, p=0.005, 1,2>3 |
| CBCL Internalizing Score (raw) | 5.18 [0-51] | 5.62 | 5.18 [0-51] | 5.58 | 4.98 [0-51] | 5.42 | F=3.47, p=0.03, 1,2>3 |
| CBCL Externalizing Score (raw) | 4.51 [0-49] | 5.88 | 4.46 [0-49] | 5.83 | 4.28 [0-49] | 5.63 | F=3.88, p=0.02, 1,2>3 |
| | **Total** | **%** | **Total** | **%** | **Total** | **%** | **X2, p-value** |
| Sex (Female) | 4369 | 47.5 | 4001 | 47.6 | 3974 | 48.8 | X2=3.46, p=0.17 |
| *Household Income* | | | | | | | X2=0.72, p=0.94 |
| <$50K | 2531 | 27.5 | 2531 | 27.5 | 2274 | 27.9 | |
| $50-$100K | 2381 | 25.9 | 2381 | 25.9 | 2080 | 25.5 | |
| >$100K | 3487 | 37.9 | 3487 | 37.9 | 3065 | 37.6 | |
| *Participant race/ethnicity* | | | | | | | X2=18.9, p=0.01 |
| White | 4704 | 51.2 | 4480 | 53.3 | 4125 | 50.7 | |
| Black | 1360 | 14.8 | 1149 | 13.6 | 1242 | 15.2 | |
| Asian | 205 | 2.2 | 176 | 2.1 | 185 | 2.27 | |
| Hispanic | 1973 | 21.5 | 1715 | 20.4 | 1764 | 21.7 | |
| Other | 948 | 10.3 | 870 | 10.3 | 815 | 10 | |
| *Parent Education* | | | | | | | X2=26.2, p<0.001 |
| <HS Diploma | 468 | 5.1 | 352 | 4.2 | 436 | 5.35 | |
| HS Diploma/GED | 887 | 9.65 | 732 | 8.71 | 810 | 9.95 | |
| Some College | 2389 | 25.9 | 2146 | 25.5 | 2082 | 25.5 | |
| Bachelors | 2287 | 24.9 | 2162 | 25.7 | 2027 | 24.9 | |
| Post-Graduate | 3150 | 34.3 | 3003 | 35.7 | 2775 | 34.1 | |

*Table S2. Singular Values and Variance Accounted for in each LV in the first analysis*

**Main Analysis (n=9191)**

| **PLS** | | **CCA** | |
|---|---|---|---|
| **Singular Values** | **Variance Explained** | **Singular Values** | **Variance Explained** |
| 0.389 | 81.6% | 0.131 | 19.32% |
| 0.107 | 6.13% | 0.112 | 14.17% |
| 0.08 | 3.49% | 0.106 | 12.68% |
| 0.066 | 2.36% | 0.098 | 10.79% |
| 0.06 | 1.95% | 0.09 | 9.15% |
| 0.049 | 1.34% | 0.084 | 7.99% |
| 0.047 | 1.17% | 0.076 | 6.59% |
| 0.036 | 0.07% | 0.073 | 6.07% |
| 0.032 | 0.05% | 0.067 | 5.06% |
| 0.025 | 0.04% | 0.064 | 4.66% |
| 0.022 | 0.02% | 0.056 | 3.48% |

*Table S3. Singular Values and Variance Accounted for in each LV in the Elevated-CBCL Sample*

| High Psychopathology (n=1016) | | | |
|---|---|---|---|
| **PLS** | | **CCA** | |
| **Singular Values** | **Variance Explained** | **Singular Values** | **Variance Explained** |
| 0.671 | 53.9% | 0.334 | 14.7% |
| 0.314 | 11.8% | 0.315 | 13.1% |
| 0.288 | 9.96% | 0.294 | 11.3% |
| 0.258 | 7.98% | 0.287 | 10.8% |
| 0.197 | 4.62% | 0.276 | 10% |
| 0.174 | 3.62% | 0.261 | 8.98% |
| 0.145 | 2.51% | 0.246 | 7.94% |
| 0.127 | 1.92% | 0.239 | 7.49% |
| 0.117 | 1.65% | 0.216 | 6.11% |
| 0.1 | 1.19% | 0.198 | 5.17% |
| 0.08 | 0.07% | 0.182 | 4.34% |

*Table S4. Singular Values and Variance Accounted for in each LV in the Symptom Endorsement Sample*

| Symptom Endorsement (n=5196) | | | |
|---|---|---|---|
| **PLS** | | **CCA** | |
| **Singular Values** | **Variance Explained** | **Singular Values** | **Variance Explained** |
| 0.38 | 59% | 0.18 | 21% |
| 0.22 | 18% | 0.14 | 13% |
| 0.13 | 6.90% | 0.14 | 12% |
| 0.11 | 4.70% | 0.13 | 10% |
| 0.09 | 3.10% | 0.12 | 9.30% |
| 0.07 | 2.10% | 0.11 | 7.80% |
| 0.06 | 1.80% | 0.1 | 6.60% |
| 0.05 | 1.20% | 0.1 | 6.20% |
| 0.05 | 0.09% | 0.09 | 5.20% |
| 0.04 | 0.06% | 0.08 | 4.40% |
| 0.03 | 0.04% | 0.08 | 3.80% |

*Table S5. Singular Values and Variance Accounted for in each LV in the second analysis*

**NIH Analysis (n=9034)**

| **PLS** | | **CCA** | |
|---|---|---|---|
| **Singular Values** | **Variance Explained** | **Singular Values** | **Variance Explained** |
| 0.429 | 75.50% | 0.205 | 41.60% |
| 0.167 | 11.40% | 0.125 | 15.50% |
| 0.124 | 6.32% | 0.115 | 13.20% |
| 0.084 | 2.88% | 0.102 | 10.50% |
| 0.064 | 1.67% | 0.091 | 8.24% |
| 0.059 | 1.45% | 0.079 | 6.15% |
| 0.042 | 0.07% | 0.069 | 4.71% |

*Table S6. Model fit of selected brain and behaviour variables when including age and age-squared as regressors.*

| Behaviour/Brain Example Variables | | | | |
|---|---|---|---|---|
| **CBCL Variables** | **Regressor** | **F-value** | **t-value** | **p-value** |
| anxiety/depression subscale | age | 0.85 | 0.92 | 0.35 |
| | age-squared | 0.61 | 0.61 | 0.54 |
| attention subscale | age | 2.32 | 1.5 | 0.12 |
| | age-squared | 1.49 | 0.8 | 0.23 |
| social problems subscale | age | 5.68 | 2.38 | 0.02 |
| | age-squared | 2.91 | 0.38 | 0.054 |
| rule-breaking subscale | age | 1.47 | 1.21 | 0.22 |
| | age-squared | 1.02 | 0.75 | 0.36 |
| **Cortical Thickness Variables** | | | | |
| left lateral OFC | age | 84.67 | 9.2 | <0.001 |
| | age-squared | 42.48 | 0.55 | 0.58 |
| left pars orbitalis | age | 50.75 | 7.12 | <0.001 |
| | age-squared | 25.25 | 0.57 | 0.56 |
| right lingual gyrus | age | 86.86 | 9.32 | <0.001 |
| | age-squared | 44.49 | 1.45 | 0.14 |

| | | | | |
|---|---|---|---|---|
| right temporal pole | age | 0.5 | 0.71 | 0.47 |
| | age-squared | 0.44 | 0.61 | 0.54 |

### References

Ameis, S.H., Ducharme, S., Albaugh, M.D., Hudziak, J.J., Botteron, K.N., Lepage, C., Zhao, L., Khundrakpam, B., Collins, D.L., Lerch, J.P., Wheeler, A., Schachar, R., Evans, A.C., Karama, S., 2014. Cortical thickness, cortico-amygdalar networks, and externalizing behaviors in healthy children. Biol Psychiatry 75, 65–72. https://doi.org/10.1016/j.biopsych.2013.06.008

Dienes, K.A., Chang, K.D., Blasey, C.M., Adleman, N.E. and Steiner, H., 2002. Characterization of children of bipolar parents by parent report CBCL. Journal of Psychiatric Research, 36(5), pp.337-345.

Gross, D., Fogg, L., Young, M., Ridge, A., Cowell, J. M., Richardson, R., & Sivan, A. (2006). The equivalence of the Child Behavior Checklist/1 1/2-5 across parent race/ethnicity, income level, and language. Psychological Assessment, 18(3), 313–323. https://doi.org/10.1037/1040-3590.18.3.313

Hall, P.A., Best, J.R., Beaton, E.A., Sakib, M.N. and Danckert, J., 2021. Morphology of the prefrontal cortex predicts body composition in early adolescence: cognitive mediators and environmental moderators in the ABCD Study. Social Cognitive and Affective Neuroscience

Hill, W.D., Hagenaars, S.P., Marioni, R.E., Harris, S.E., Liewald, D.C.M., Davies, G., Okbay, A., McIntosh, A.M., Gale, C.R., Deary, I.J., 2016. Molecular Genetic Contributions to Social Deprivation and Household Income in UK Biobank. Current Biology 26, 3083–3089. https://doi.org/10.1016/j.cub.2016.09.035

Lawson, G.M., Duda, J.T., Avants, B.B., Wu, J., Farah, M.J., 2013. Associations between children's socioeconomic status and prefrontal cortical thickness. Dev Sci 16, 641–652. https://doi.org/10.1111/desc.12096

McIntosh, A.R., Lobaugh, N.J., 2004. Partial least squares analysis of neuroimaging data: Applications and advances. Neuroimage 23, 250–263. https://doi.org/10.1016/j.neuroimage.2004.07.020

Modabbernia, A., Janiri, D., Doucet, G.E., Reichenberg, A., Frangou, S., 2021. Multivariate Patterns of Brain-Behavior-Environment Associations in the Adolescent Brain and Cognitive Development Study. Biol Psychiatry 89, 510–520. https://doi.org/10.1016/j.biopsych.2020.08.014

Myers, L. and Sirois, M.J., 2006. Spearman correlation coefficients, differences between. Encyclopedia of statistical sciences, 12. https://doi.org/10.1002/0471667196.ess5050.pub2

Owens, M.M., Potter, A., Hyatt, C.S., Albaugh, M., Thompson, W.K., Jernigan, T., Yuan, D., Hahn, S., Allgaier, N. and Garavan, H., 2021. Recalibrating expectations about effect size: A multi-method survey of effect sizes in the ABCD study. PloS one 16, p.e0257535. https://doi.org/10.1371/journal.pone.0257535

Piccolo, L.R., Merz, E.C., He, X., Sowell, E.R., Noble, K.G., 2016. Age-related differences in cortical thickness vary by socioeconomic status. PLoS One 11, 1–18. https://doi.org/10.1371/journal.pone.0162511

Rakesh, D., Zalesky, A., Whittle, S., 2021. Similar but distinct – Effects of different socioeconomic indicators on resting state functional connectivity: Findings from the Adolescent Brain Cognitive Development (ABCD) Study®. Dev Cogn Neurosci 51, 101005. https://doi.org/10.1016/j.dcn.2021.101005

Tollenaar, M.S., Beijers, R., Garg, E., Nguyen, T.T., Lin, D.T., MacIsaac, J.L., Shalev, I., Kobor, M.S., Meaney, M.J., O'Donnell, K.J. and de Weerth, C., 2021. Internalizing symptoms associate with the pace of epigenetic aging in childhood. Biological Psychology, 159, p.108021.

Wilde, E.A., Merkley, T.L., Bigler, E.D., Max, J.E., Schmidt, A.T., Ayoub, K.W., McCauley, S.R., Hunter, J. V., Hanten, G., Li, X., Chu, Z.D., Levin, H.S., 2012. Longitudinal changes in cortical thickness in children after traumatic brain injury and their relation to behavioral regulation and emotional control. International Journal of Developmental Neuroscience 30, 267–276. https://doi.org/10.1016/j.ijdevneu.2012.01.003

Zhu, X., Ward, J., Cullen, B. et al. Phenotypic and genetic associations between anhedonia and brain structure in UK Biobank. Transl Psychiatry 11, 395 (2021). https://doi.org/10.1038/s41398-021-01522-4
